# Syndapin and GTPase RAP-1 Control Endocytic Recycling via RHO-1 and Non-Muscle Myosin II

**DOI:** 10.1101/2023.02.27.530328

**Authors:** Wilmer R. Rodriguez-Polanco, Anne Norris, Agustin B. Velasco, Adenrele M. Gleason, Barth D. Grant

## Abstract

After endocytosis, many plasma membrane components are recycled via narrow-diameter membrane tubules that emerge from early endosomes to form recycling endosomes, eventually leading to their return to the plasma membrane. We previously showed that the F-BAR and SH3 domain Syndapin/PACSIN-family protein SDPN-1 is required *in vivo* for basolateral endocytic recycling in the *C. elegans* intestine. Here we sought to determine the significance of a predicted interaction between the SDPN-1 SH3 domain and a target sequence in PXF-1/PDZ- GEF1/RAPGEF2, a known exchange factor for Rap-GTPases. We found that endogenous mutations we engineered into the SDPN-1 SH3 domain, or its binding site in the PXF-1 protein, interfere with recycling *in vivo*, as does loss of the PXF-1 target RAP-1. Rap-GTPases have been shown in several contexts to negatively regulate RhoA activity. Our results show that RHO- 1/RhoA is enriched on SDPN-1 and RAP-1 positive endosomes in the *C. elegans* intestine, and loss of SDPN-1 or RAP-1 elevates RHO-1(GTP) levels on intestinal endosomes. Furthermore, we found that depletion of RHO-1 suppressed *sdpn-1* mutant recycling defects, indicating that control of RHO-1 activity is a key mechanism by which SDPN-1 acts to promote endocytic recycling. RHO-1/RhoA is well-known for controlling actomyosin contraction cycles, although little is known of non-muscle myosin II on endosomes. Our analysis found that non-muscle myosin II is enriched on SDPN-1 positive endosomes, with two non-muscle myosin II heavy chain isoforms acting in apparent opposition. Depletion of *nmy-2* inhibited recycling like *sdpn-1* mutants, while depletion of *nmy-1* suppressed *sdpn-1* mutant recycling defects, indicating actomyosin contractility in controlling recycling endosome function.

## Introduction

Cells internalize extracellular material, receptor-associated ligands, membrane proteins and lipids by endocytosis (1). Uptake mechanisms can be generally divided into clathrin-dependent and clathrin-independent endocytosis (CDE and CIE, respectively), after which internalized cargo is sorted within endosomes for delivery to distinct intracellular destinations (2). As membrane and fluid are added to the early endosome by incoming vesicles, narrow-diameter recycling tubules form, collecting a subset of cargo (3). Most plasma membrane lipids, and many types of internalized transmembrane proteins, are recycled back to the plasma membrane either directly (rapid recycling), indirectly via the endocytic recycling compartment (ERC) (slow recycling), or Golgi (retrograde recycling) (3, 4). As early endosomes mature they become late endosomes, eventually fusing with lysosomes, promoting degradation of remaining cargo. Many cellular processes depend upon accurate and efficient endocytic recycling, including cell migration, cell division, cell-cell fusion events, and control of neuronal signaling during learning and memory (3, 4). Trafficking pathways are often more elaborate in the context of polarized cells such as epithelia and neurons that maintain multiple functionally distinct plasma membrane domains, e.g. basolateral versus apical in epithelia, and dendrite versus axon in neurons (4).

The *C. elegans* intestine is a polarized epithelial tube consisting of exactly 20 cells that form nine donut-like rings, the most anterior ring consisting of four symmetrical cells and all other rings comprised of two cells (5, 6). The intestinal cells have a basolateral surface that faces the pseudocoelom and controls the exchange of molecules between the intestine and the rest of the body (5). The microvilli-rich apical surface faces the central lumen and is important for nutrient uptake (6). We previously developed the *C. elegans* intestine as a model system for the study of polarized epithelial trafficking mechanisms, allowing analysis of transport in the native context of a polarized-epithelial tube within the intact living animal (7–10). Taking advantage of the *C. elegans* intestinal system, our laboratory previously showed that *C. elegans* SDPN-1, a Syndapin/PACSIN family protein, is important for basolateral endocytic recycling *in vivo,* a finding confirmed independently by others (9, 11).

Syndapin/PACSIN family proteins bear a single N-terminal F-BAR domain and a single C-terminal SH3 domain. BAR domains in general, including the F-BAR subclass, are known to sense membrane curvature and to promote positive curvature, properties likely to help drive membrane budding and tubulation (12). Our previous work showed that *C. elegans* SDPN-1 is essential for the basolateral recycling of cargo proteins that are transported via the early endosome–to–recycling endosome route, but not other routes (Gleason et al., 2016). Affected cargos accumulate at much higher-than-normal levels in the endosomes of *sdpn-1* mutants (Gleason et al., 2016). These include well characterized model cargo proteins hTAC (human IL-2 receptor α-chain), hTFR (human transferrin receptor), and DAF-4 (type-II TGF-β receptor), but not MIG-14/Wls or SMA-6 (type-I TGF-β receptor) that recycle via the retromer pathway, or CD4-LL that does not recycle (Gleason et al., 2016). SDPN-1 protein is highly enriched on early and basolateral recycling endosomes of the intestinal cells, and *sdpn-1 null* mutants display abnormal accumulation of early and basolateral recycling endosomes, suggesting a direct role in their regulation (Gleason et al. 2016). Mammalian Syndapin2 is also well known to associate with recycling endosome proteins and to promote endocytic recycling, but many questions remain about the mechanism by which Syndapin/PACSIN proteins regulate endosomal function (13–15).

To better understand how SDPN-1 promotes recycling we sought to identify partner proteins relevant to recycling, with a particular focus on results from the *C. elegans* SH3-ome project (16). Here we investigated one high-confidence hit for the SDPN-1 SH3 domain, PXF-1/PDZ-GEF, an exchange factor for the small Ras-like GTPase RAP-1/Rap (17, 18). Importantly we show that PXF-1 and its target RAP-1 also regulate recycling like SDPN-1, and interfering with the interaction of SDPN-1 and PXF-1 disrupts recycling. We trace the activity of RAP-1 on endosomes to negative regulation of RHO-1 and myosin II heavy chain isoform NMY-1, and show that reduction in RHO-1 or NMY-1 greatly ameliorates *sdpn-1* mutant trafficking defects. This function appears to work in opposition to myosin II heavy chain isoform NMY-2 that we find is required for recycling. Taken together our results indicate that myosin II regulation is a key output of endosomal SDPN-1 that is essential to maintain recycling function.

## Materials and Methods

### General methods and strains

All *C. elegans* strains were derived from Bristol strain N2 and cultured at 20°C on NGM plates seeded with the *Escherichia coli* strain OP50. A list of strains used in this study can be found in Supplemental Table 1.

### Transgenic Strains

To construct GFP or RFP fusion transgenes for expression in the worm intestine, Gateway destination vectors were used. Each vector contained the promoter region of the intestine-specific gene *vha-6*, followed by GFP, mScarlet, or tagRFP coding sequences, a Gateway cassette (Invitrogen, Carlsbad, CA), and *let-858* 3’ UTR sequences. The genomic or cDNA sequences of *C. elegans* genes were cloned individually into entry vector pDONR221. Mutant forms of the coding regions were obtained using site-directed mutagenesis using the Quikchange II kit (Stratagene). Coding regions were transferred into P*vha-6*-GFP or P*vha-6*-RFP vectors by LR reaction (Invitrogen). Constructs were bombarded into *unc-119(ed3)* mutant animals to establish low copy integrated transgenic lines, or integrated in single copy using the MiniMos system and Hygromycin or G418 selection (19).

### RNAi

*C. elegans* RNAi-mediated interference was performed as described (20). Feeding constructs in this study were from the Ahringer library, except for *rho-1*, for which a cDNA was cloned into the L4440/pPD129.36 vector between unique SacI and HindIII sites (21). RNAi clones for *nmy-1* and *nmy-2* were kindly provided by Adriana Dawes (Ohio State University) (22). For most RNAi experiments, synchronized L4 stage animals were grown on RNAi plates at 20°C. After the first generation, we transferred about 60 L4 stage animals to fresh RNAi plates at 20°C for another 24 hours prior to imaging as young adults.

### Protein Expression, Pulldown Assays, and Western Analysis

N-terminally hemagglutinin (HA)-tagged PXF-1 protein, amino acids 1-477, with or without mutation of the predicted SDPN-1 binding site, was synthesized *in vitro* using the TNT-coupled transcription-translation system (Promega, Madison, WI) using DNA templates pcDNA3.1– 2xHA-PXF-1 and pcDNA3.1–2xHA-PXF-1(K191/P195/P198->AAA), respectively (1.6 ug/each 50- ul reaction). The reaction cocktail was incubated at 30°C for 90 min. Control glutathione *S*- transferase (pGEX-2T), GST-SDPN-1 SH3 (final 112 amino acids), and GST-SDPN-1 SH3 (P479L) fusion proteins were expressed in the *ArcticExpress* strain of *Escherichia coli* (Stratagene, La Jolla, CA). Bacterial pellets were lysed in 10 ml B-PER Bacterial Protein Extraction Reagent (Pierce, Rockford, ILL) with Complete Protease Inhibitor Cocktail Tablets (Roche, Indianapolis, IN). Extracts were cleared by centrifugation, and supernatants were incubated with glutathione-Sepharose 4B beads (Amersham Pharmacia, Piscataway, NJ) at 4°C overnight.

Beads were then washed six times with cold STET buffer (10 mM Tris-HCl, pH 8.0, 150 mM NaCl, 1 mM EDTA, 0.1% Tween-20). In vitro–synthesized HA-tagged protein (10 ul TNT mix diluted in 500 ul STET) was added to the beads and allowed to bind at 4°C for 2 h. After six additional washes in STET the proteins were eluted by boiling in 30 ul 2X-SDS-PAGE sample buffer. Eluted proteins were separated on SDS-PAGE, blotted to nitrocellulose, visualized with Ponceau S and scanned, then washed to remove Ponceau staining, and probed with anti-HA (16B12) antibodies and HA-conjugated secondary antibodies (Pierce). Detection was achieved using Super Signal West Pico detection system (Pierce).

### CRISPR

*sdpn-1* P479L and *pxf-1* K191/P195/P198->AAA mutations were introduced into the endogenous genes in *C. elegans* using the *dpy-10* co-CRISPR technique as described by Paix and Seydoux (23), creating mutant alleles *sdpn-1(pw33)* and *pxf-1(pw34)*. 3XFLAG-GFP was inserted into the *pxf-1* gene between the last codon and stop codon using the SEC selection technique described by Dickinson as modified by Schwartz (24, 25).

### Microscopy and image analysis

Imaging was performed on live first-day adult hermaphrodites animals mounted on slides using 5% agarose pads with 10 mM levamisole. Most single channel data in Figure 2 was captured on a Zeiss LSM 710 confocal microscope. All other single channel data, including **Figure 2B**-**B’,** was captured on a Zeiss LSM 800 confocal microscope. For both microscopes, the LSM 710 and LSM 800, GFP tagged protein image acquisition used the 488 nm laser line and the 63X (1.4 NA) objective and ZEISS ZEN 2.3 imaging software. True GFP signal was separated from the broad- spectrum auto-fluorescence displayed by the lysosome-related organelles (gut granules) of the *C. elegans* intestine using the Spectral Fingerprinting and Linear Unmixing functions in ZEN. The images were exported as TIF files for analysis in Metamorph version 7.7 software.

The “Integrated Morphometry Analysis” function of Metamorph was used to measure fluorescence intensity, puncta number (structure count), and fluorescence area (total area) within unit regions. At least 9 animals were used for each analysis. Fluorescence signal in the basolateral membrane between two neighboring cells was avoided for the analysis.

For multiwavelength fluorescence colocalization studies, images were obtained using a spinning-disk confocal imaging system: Axiovert Z1 microscope (Carl Zeiss Microimaging) equipped with a X-Light V2 Spinning Disk Confocal Unit (CrestOptics), 7-line LDI Laser Launch (89 North), Prime 95B Scientific CMOS camera (Photometrics) and 63X and 100X oil-immersion objectives. Data were captured as an image stack with 0.2 μm step size using Metamorph version 7.7 software (Universal Imaging), and deconvolved using Autoquant X3 (Media Cybernetics). In Metamorph, a single plane from each image stack (either top plane or middle plane) was chosen in which both markers were clearly visible. In the middle plane of the worm intestine, the apical membranes and the basolateral membranes are visible, while on top plane images only the basolateral membrane is visible. Images taken in the DAPI channel were used to identify broad-spectrum intestinal autofluorescence caused by lysosome-like organelles. The DAPI channel image was subtracted from the green and red channel images to remove autofluorescent signal from the colocalization analysis. The DAPI subtraction was done using the Metamorph “Arithmetic” function at 16-bit depth. Quantification of colocalization (Manders coefficients) in autofluorescence subtracted images was performed using open source Fiji (Image J) software and the “Colocalization Threshold” function. At least 9 animals were used for each analysis. For middle plane images, the fluorescence signal in the apical membrane and the basolateral membrane between two neighboring cells were avoided.

### Statistical analysis

Quantified data obtained in this study was graphed and statistically analyzed using GraphPad Prism 7.04 software.

## Results

### The SDPN-1 SH3 domain binds to PXF-1 via a specific proline-rich sequence

Syndapin family proteins, also known as PACSINs, contain an N-terminal F-BAR domain (FCH- BAR, Fes/CIP4 homology-Bin/Amphiphysin/Rvs) and a C-terminal SH3 (Src-homology 3) domain (14). *C. elegans* has a single Syndapin family member SDPN-1, while the situation in mammals is more complex, with three Syndapin family members Syndapin1/PACSIN1, Syndapin2/PACSIN2, and Syndapin3/PACSIN3 (26). Syndapins form dimers through their F-BAR domains, with the F- BAR dimer forming a mildly curved structure that binds directly to membrane phospholipids (14). Higher order assemblies of these proteins scaffold the membrane to promote bending and remodeling, in many cases producing membrane tubulation (26). Indeed, our previous work showed that full-length recombinant *C. elegans* SDPN-1 tubulates acidic membranes *in vitro* (9). *In vivo* SDPN-1 localizes to early endosomes and basolateral recycling endosomes in the *C. elegans* intestine, and is required for basolateral recycling, but the molecular mechanism(s) by which it mediates endocytic recycling remain poorly understood (9). To gain a better understanding of how SDPN-1 promotes endosomal transport we sought to identify partner proteins that work with SDPN-1 in this process.

The SH3 domain is a peptide recognition module for proline-rich sequences that is well-known for mediating specific protein-protein interactions (27). Thus, we focused on the SDPN-1 SH3 domain as a molecular entry point to identify direct SDPN-1 binding partners. We were aided in this goal by the *C. elegans* SH3-interactome project (16), a large-scale analysis that used peptide phage display to define the binding preferences for most SH3-domains encoded in the *C. elegans* genome, including generating a weighted SDPN-1 SH3 consensus target sequence (Fig. 1A). The SH3 interactome project also screened directly for SDPN-1 SH3-domain interacting proteins using a stringent yeast-2-hybrid approach. Importantly, both approaches identified the protein PXF-1 as a prime SDPN-1 SH3 domain candidate. The project recovered PXF-1 via genome-wide yeast-2-hybrid with a SDPN-1 SH3 bait, but not any of the other SH3 domain baits, and independently identified PXF-1 amino acids 191-198 as the second-best match for the SDPN-1 SH3-binding consensus in the entire predicted *C. elegans* proteome (16).

**Figure 1.**
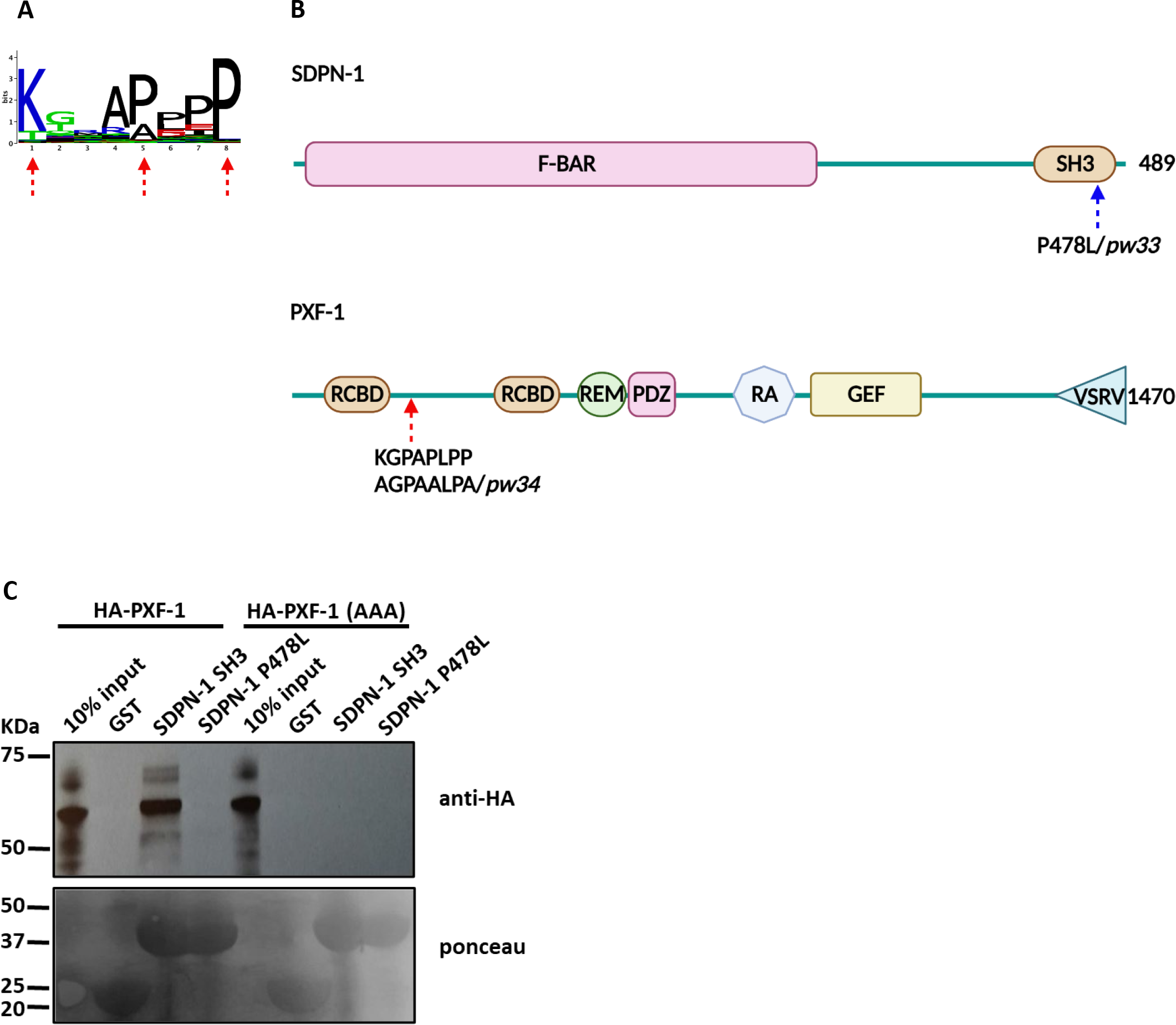
The SDPN-1 SH3 domain binds to PXF- 1. (A) SDPN-1 SH3 domain consensus target sequence; red arrows indicate conserved residues that were mutated to alanine (*pxf-1(pw34)*) to disrupt interaction with the SDPN-1 SH3 domain. **(B)** Schematic representation of SDPN-1 and PXF-1 domain structure from the N-terminal to the C-terminal; the blue arrow indicates the *sdpn-1(pw33)* mutation site and the red arrow indicates the *pxf- 1(pw34)* mutation site. **(C)** Glutathione beads loaded with recombinant GST or the GST-SDPN-1 SH3 domain were incubated with *in vitro* translated HA-tagged PXF-1 *wild-type* or aa191-198 (**K**GPA**P**LP**P** to **A**GPA**A**LP**A**). Bound proteins were eluted and analyzed by Western blot probed with anti-HA antibody. The *wild-type* HA-tagged PXF-1 was bound by *wild-type* GST-SDPN-1-SH3, but not by mutant (P478L), or unfused GST. The (KGPAPLPP to AGPAALPA) mutant HA-tagged PXF-1 failed to bind GST alone or GST-SDPN-1-SH3 *wild-type* and mutant domains. Total GST and GST-SDPN-1-SH3 bait proteins were visualized by Ponceau Staining prior to antibody staining.

PXF-1 (PDZ-GEF or RAGEF in mammals) is a well-characterized exchange factor for the small Ras-like GTPases RAP-1/Rap1 and RAP-2/Rap2, and Rap-GTPases have been implicated in some endocytic recycling events in other organisms, further suggesting that a SDPN-1/PXF-1 interaction could be relevant to understanding the role of SDPN-1 in recycling (17, 28–30). PXF-1 has a complex domain structure, with two RCBD-like domains: one near the N-terminus, and the other more centrally located(17, 31). While RCBD domains in some proteins bind to cAMP, biochemical analysis of human PDZ-GEF2, a PXF-1 ortholog, failed to identify any cyclic- nucleotide binding for these RCBD-like domains (32). The predicted SDPN-1 SH3 binding site at residues 191-198 resides in a predicted unstructured region between the two RCBD-like domains (https://alphafold.ebi.ac.uk/entry/G5EDB9). PXF-1 also contains REM (Ras Exchange Motif), PDZ (PSD-95, Dlg, and ZO-1 homology), RA (Ras Association), and GEF (Guanine nucleotide Exchange Factor) domains, as well as a C-terminal VSRV sequence that may interact with PDZ-domains (Fig 1B) (17).

To better analyze the ability of the SDPN-1 SH3 domain to bind to PXF-1, and test the importance of the predicted binding site identified as a strong match to the phage-display consensus sequence, we performed GST pulldown experiments. As baits we compared GST, GST-SH3^sdpn-1^(+), and GST-SH3^sdpn-1^(P478L). The P478L mutation in the SDPN-1 SH3 domain is equivalent to mammalian Syndapin I P434L, which was previously shown to abolish the ability of the Syndapin I SH3 domain to bind partner proteins (33). These baits were paired with *in vitro* translated HA-PXF-1(1–478) bearing the wild-type sequence, or with the consensus- matching PXF-1(191–198) SH3 binding site mutated (**K**GPA**P**LP**P** to **A**GPA**A**LP**A**) (Fig 1A-B). We found that HA-PXF-1(+) interacted robustly with GST-SH3^sdpn-1^(+), but not GST or GST-SH3^sdpn- 1^(P478L), indicating the specificity of the interaction. Moreover, we found that mutation of HA- PXF-1 at the predicted interaction sequence (191–198) abolished the interaction with GST- SH3^sdpn-1^(+), despite other proline-rich sequences remaining in the PXF-1 prey protein. Taken together these results indicate that the SDPN-1 SH3 domain can bind to PXF-1, and the predicted binding site at aa191-198 is crucial for this interaction.

### The SDPN-1/PXF-1 interaction is important for endocytic recycling

Given our binding results we hypothesized that the interaction of the SDPN-1 SH3-domain with PXF-1 is important for SDPN-1-mediated regulation of endocytic recycling. To interfere with this interaction *in vivo* we used CRISPR-mediated genome editing to create *C. elegans* mutants *sdpn-1(pw33)* and *pxf-1(pw34)*, in which the same point mutations that interfered with the SDPN-1/PXF-1 binding interaction *in vitro* were engineered into the endogenous *sdpn-1* and *pxf-1* genes *in vivo* (Fig. 1b). While *pxf-1* null mutations are lethal (17), *pxf-1(pw34)* is a viable allele.

In particular we found that the interaction-defective mutants *sdpn-1(pw33)* and *pxf-1(pw34)* displayed gross over-accumulation of intestinal GFP::RME-1 marked basolateral recycling endosomes and GFP::RAB-5 marked early endosomes in the central cytoplasm (middle plane Fig. 2K), very similar to phenotypes we had previously observed in null mutants for *sdpn-1* (Fig. 2, panel A-A’’ and B-B’’; Fig. S1 A, A”) (9). For GFP::RME-1 labeled recycling endosomes we measured endosome fluorescence intensity and number. In the case of GFP::RAB-5, the mutants accumulated so many early endosomal compartments that it was not possible to segment the images to quantify endosome number, so in addition to fluorescence intensity we measured the the percent of total cytoplasmic area labeled by GFP::RAB-5.

**Figure 2.**
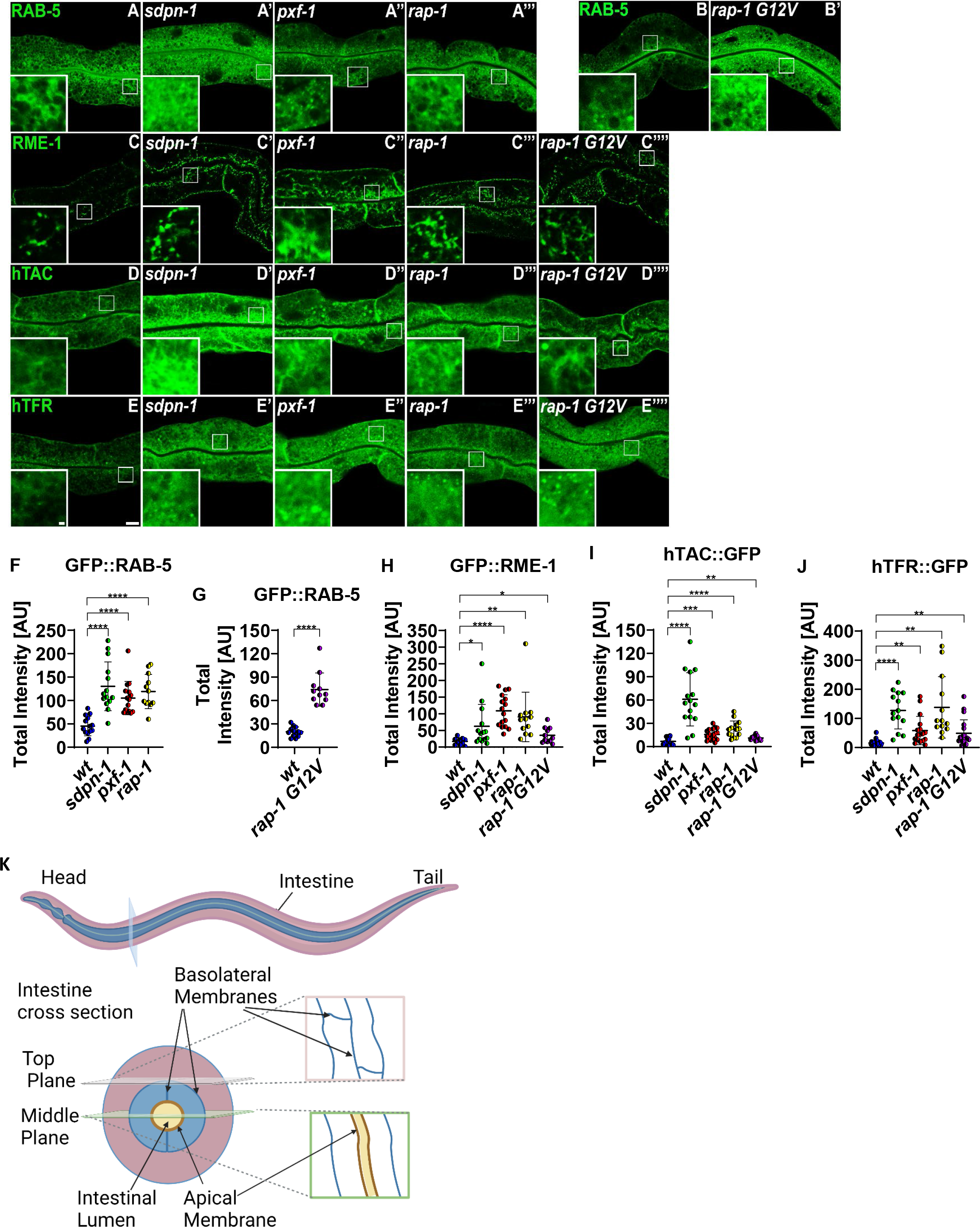
SDPN-1 interaction with PXF-1, and RAP-1 is important for proper endocytic recycling. All images were obtained by confocal laser-scanning microscopy in intact living animals expressing intestine-specific GFP-tagged proteins. Representative confocal images of GFP::RAB-5 expressed in a *wild-type* **(A)**, *sdpn-1(pw33)* **(A’)**, *pxf-1(pw34)* **(A’’)**, and *rap-1(0)* **(A’’’)** background are shown. SDPN-1 and PXF-1 interacting-defective mutations, and loss of RAP-1, causes intracellular accumulation of compartments labeled by early endosome marker GFP::RAB-5 when compared with wild-type. In addition, GFP::RAB-5 expressed in a wild-type **(B)**, and *rap- 1(re180)* GTP-locked mutant **(B’)** background also showed an increase in the intracellular compartments labeled by early endosome marker GFP::RAB-5. Confocal images of GFP::RME-1 basolateral recycling endosome marker expressed in a *wild-type* **(C)**, *sdpn-1(pw33)* **(C’)**, *pxf-1(pw34)* **(C’’)**, *rap-1(0)* **(C’’’)**, and *rap-1(re180)* **(C’’’’)** background are shown. There is an increase of intracellular recycling compartments signal when SDPN-1 and PXF-1 interaction is lost, and when RAP-1 is missing or is GTP-locked. Micrographs showing representative confocal images of model GFP-tagged recycling cargo protein, the human interleukin-2 receptor (hTAC::GFP), expressed in a *wild-type* **(D)**, *sdpn-1(pw33)* **(D’)**, *pxf-1(pw34)* **(D’’)**, *rap-1(0)* **(D’’’)**, and *rap-1(re180)* **(D’’’’)** background. Micrographs showing representative confocal images of another model GFP-tagged recycling cargo protein, the human transferrin receptor (hTFR::GFP), expressed in a *wild-type* **(E)**, *sdpn-1(pw33)* **(E’)**, *pxf- 1(pw34)* **(E’’)**, *rap-1(0)* **(E’’’)**, and *rap-1(re180)* **(E’’’’)** background. Losing the interaction between SDPN-1 and PXF-1, and when RAP-1 is missing or is GTP-locked, displays a significant increase in the intracellular accumulation of both recycling cargoes hTAC::GFP and hTFR::GFP. Main panel scale bars = 10 µm. Magnified inset panel scale bars = 1 µm. **(F)** Quantification of total fluorescence intensity for **(A-A’’’)**. **(G)** Quantification of total fluorescence intensity for **(B-B’)**. **(H)** Quantification of total fluorescence intensity for **(C-C’’’’)**. **(I)** Quantification of total fluorescence intensity for **(D-D’’’’)**. **(J)** Quantification of total fluorescence intensity for **(E-E’’’’)**. **(K)** *C. elegans* intestinal diagram. Each data point represents one animal. Error bars represent SEM: *p< 0.05, **p< 0.01, *** p< 0.001, **** p< 0.0001 (Two-tailed Welch’s *t* test).

We also tested the importance of the SDPN-1/PXF-1 interaction in cargo trafficking, focused on well-characterized model endocytic recycling cargoes markers hTAC (human IL2-receptor alpha- chain) and hTFR (human transferrin receptor) tagged with GFP, both of which depend upon SDPN-1 for their recycling (9, 10). hTAC is a clathrin-independent cargo that recycles via the recycling endosome in an ARF-6-dependent pathway (34, 35), while hTFR is clathrin-dependent in its endocytosis and also recycles via recycling endosomes (10). Previous studies showed that *sdpn-1* null mutants display dramatic endosomal accumulation of hTAC and hTFR within the intestinal cells (9). Importantly, we observed that the interaction-defective mutants caused intracellular accumulation of recycling cargoes similar to that previously observed upon the complete loss of SDPN-1 (Fig. 2, panel C-C’’ and D-D’’). Taken together, our results indicate that a SDPN-1/PXF-1 interaction is important for the function of the basolateral recycling pathway.

### The RAP-1 GTPase cycle is required for endocytic recycling

Since *C. elegans* PXF-1 and its mammalian homologs are specific GDP/GTP exchange factors for Rap-GTPases *in vivo* and *in vitro* (17, 36, 37), we surmised that the role of the SDPN-1/PXF-1 interaction is to activate Rap-GTPases to promote endocytic recycling. To test this hypothesis, we analyzed the effect of a *rap-1*(*0*) mutant on relevant markers *in vivo*. Indeed, we found that *rap-1* null mutants displayed significantly increased accumulation of RAB-5 marked early endosomes and RME-1 marked recycling endosomes, as well as increased intracellular accumulation of recycling cargo markers hTAC::GFP and hTFR::GFP, supporting this hypothesis (Fig. 2, panel A’’’, B’’’, C’’’, D’’’, F, H-J; Fig. S1 A, A”). Interestingly, we also identified abnormal accumulation defects for recycling cargo and endosome markers in a GTPase-defective *rap- 1(re180*) mutant (38), bearing the G12V mutation at the endogenous gene (Fig. 2B’,C’’’’, D’’’’, E’’’’, F, G, I-J; Fig. S1 A’). These results indicate a requirement for the full RAP-1 GTP/GDP-cycle, rather than a simple requirement for active RAP-1. We also tested for effects of *sdpn-1*(*0*) and *rap-1*(*0*) mutants on GFP-tagged *C. elegans* GLUT1 homolog FGT-1, a glucose transporter known to be naturally expressed in the *C. elegans* intestine (39). Loss of SDPN-1 and RAP-1 produced similar intracellular retention of FGT-1::GFP, also suggesting a defect in recycling (Fig. S2). Taken together these results support an important role for RAP-1 in regulating recycling downstream of SDPN-1 and PXF-1.

### PXF-1 and RAP-1 are enriched on early and basolateral recycling endosomes

To further interrogate SDPN-1, PXF-1, and RAP-1 for shared function together in recycling, we compared the localization of PXF-1 and RAP-1 with SDPN-1 in the context of the intestinal epithelium within the intact whole animal. We also compared RAP-1 localization to independent early endosome and basolateral recycling endosome markers with which SDPN-1 is known to colocalize (9). We found that endogenously tagged PXF-1::GFP colocalized with intestinal SDPN-1::tagRFP indicating significant residence of PXF-1 with SDPN-1 on basolateral endosomes (Fig 3 A-A’’’’and E). We also identified strong colocalization of intestinal SDPN- 1::GFP with tagRFP::RAP-1, indicating that at any given time most SDPN-1 overlaps with RAP-1 on the same endosomes (Fig 3 B-B’’’’and E). We did note that RAP-1 distribution appeared broader than that of SDPN-1, as reflected in a lesser but still substantial Manders’ coefficient for the reverse comparison (Fig 3E). RAP-1 labeling of structures that were SDPN-1-negative could reflect endosomal microdomains lacking SDPN-1, and/or additional RAP-1 positive organelle types such as late endosomes (40). We also observed considerable overlap between tagRFP::RAP-1 and basolateral recycling endosome marker GFP::RME-1 (Fig 3C-C’’’), and early endosome marker GFP::RAB-5 (Fig 3D-D’’’), further supporting the interpretation that SDPN-1 and RAP-1 are present on the same early and basolateral recycling endosomes where they could function together to regulate endocytic recycling.

**Figure 3.**
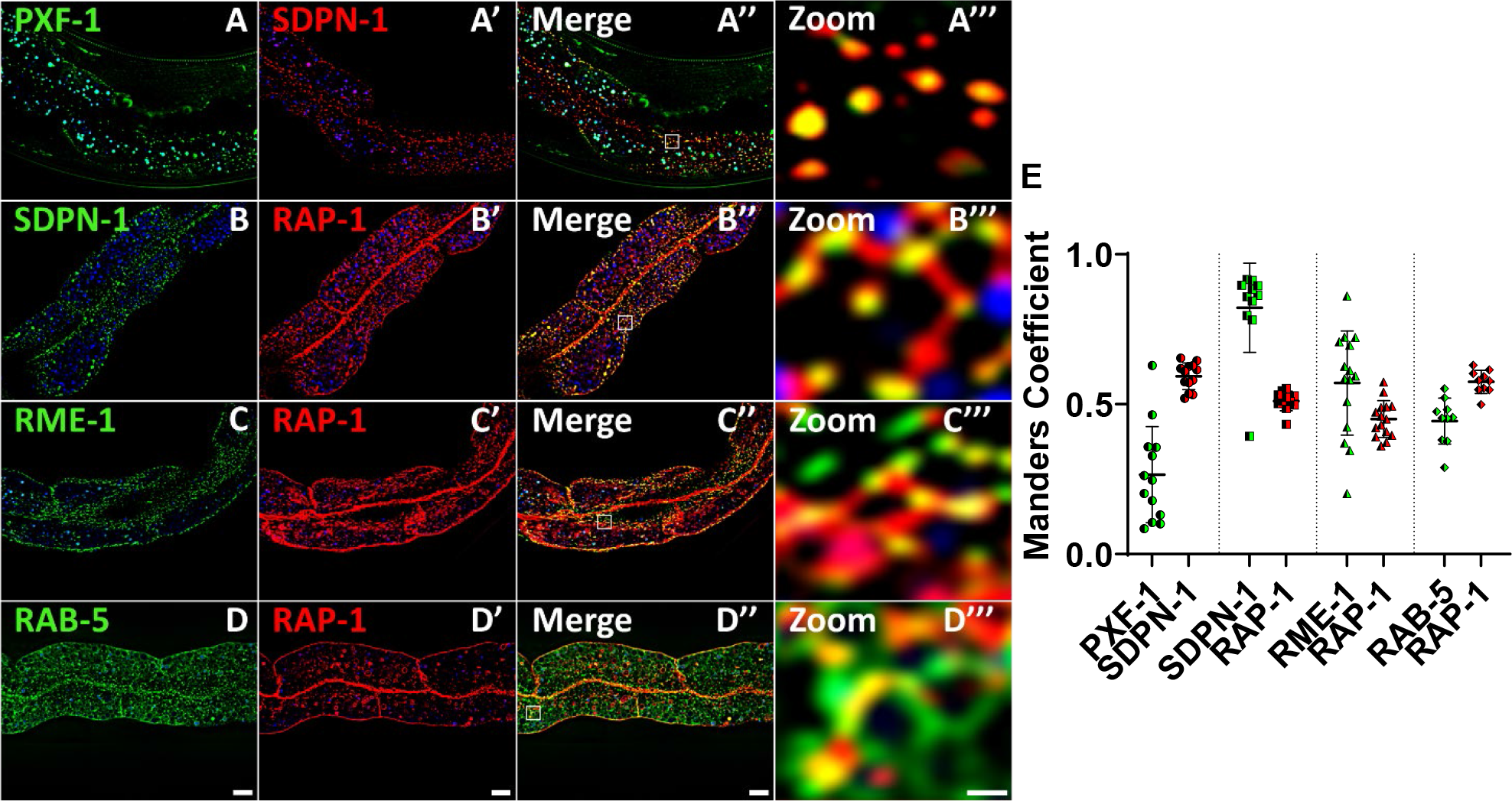
**PXF-1 and RAP-1 are enriched on SDPN-1 positive endosomes**. All micrographs are from three- dimensional (3D) confocal image stacks acquired in intact living animals expressing intestine-specific GFP and RFP-tagged proteins. **(A-A’’)** GFP::PXF-1 is found on SDPN-1 positive compartments. **(A’’’)** Magnified image from the section designated by the white rectangle in panel A’’. **(B-B’’)** tagRFP::RAP-1 colocalizes with SDPN-1 positive endosomes. **(B’’’)** Magnified image from the section designated by the white rectangle in panel B’’. **(C-C’’)** tagRFP::RAP-1 is present in GFP::RME-1-labeled basolateral recycling endosomes. **(C’’’)** Magnified image from the section designated by the white rectangle in panel C’’. **(D-D’’)** There is colocalization between tagRFP::RAP- 1 and GFP::RAB-5-labeled early endosomes. Scale bar = 10 µm. **(D’’’)** Magnified image from the section designated by the white rectangle in panel D’’. Scale bar = 1 µm. In each image, the blue channel denotes autofluorescent lysosome-like organelles, whereas the green channel denotes GFP and the red channel denotes RFP. The green channel and red channel signal that do not overlap with the blue channel signal are considered *bona fide* GFP or RFP signals respectively. **(E)** Manders coefficient for colocalization of GFP::PXF-1 with tagRFP::SDPN-1, and tagRFP::RAP-1 with SDPN-1::GFP, GFP::RME-1 and GFP::RAB-5. Each data point represents one animal.

### RHO-1 is enriched on SDPN-1 and RAP-1 positive endosomes

Given our findings indicating that RAP-1 signaling functions downstream of SDPN-1 to promote recycling, we sought to better understand the relevant target of RAP-1 for this process. While previous studies of Rap-GTPases had not identified a target of Rap-signaling for regulation of recycling, some reports indicated that mammalian Rap1 or Rap2 can negatively regulate RhoA to promote mammalian epithelial cell spreading, endothelial barrier function, or ECM stiffness mechanotransduction, in part via ARHGAP29 (41–45). We hypothesized that misregulation of endosomal Rho-GTPase activity could explain *sdpn-1/pxf-1/rap-1* associated phenotypes.

Supporting this notion, we found that mutation of *spv-1*, the closest *C. elegans* homolog of ARFGAP29, resulted in an increase in RME-1 labeled recycling endosome number, similar to the defect displayed in *pxf-1* and *rap-1* mutants (Fig. S3).

Given these results, we tested for a role of SDPN-1 and RAP-1 in the regulation of endosomal RHO-1, the only *C. elegans* homolog of RhoA (46). First, we assayed for enrichment of RHO-1 on the relevant endosomes in the *C. elegans* intestine, the expected location for direct function as part of the basolateral recycling pathway. Indeed we found that nearly all SDPN-1::GFP positive structures contained tagRFP::RHO-1 (Fig 4 A-A’’’’ and 4C), while tagRFP::RHO-1 was also present on SDPN-1 negative structures (Fig 4 A-A’’’’ and 4C). Importantly, GFP::RHO-1 and tagRFP::RAP- 1 colocalized very well, with most endosomes positive for one also positive for the other (Fig.4 B-B’’’ and 4C). Taken together, these results indicate that RHO-1 is enriched on the same endosomes as SDPN-1 and RAP-1, pointing to the potential for a SDPN-1/PXF-1/RAP-1 module influencing endosomal RHO-1 activity.

**Figure 4.**
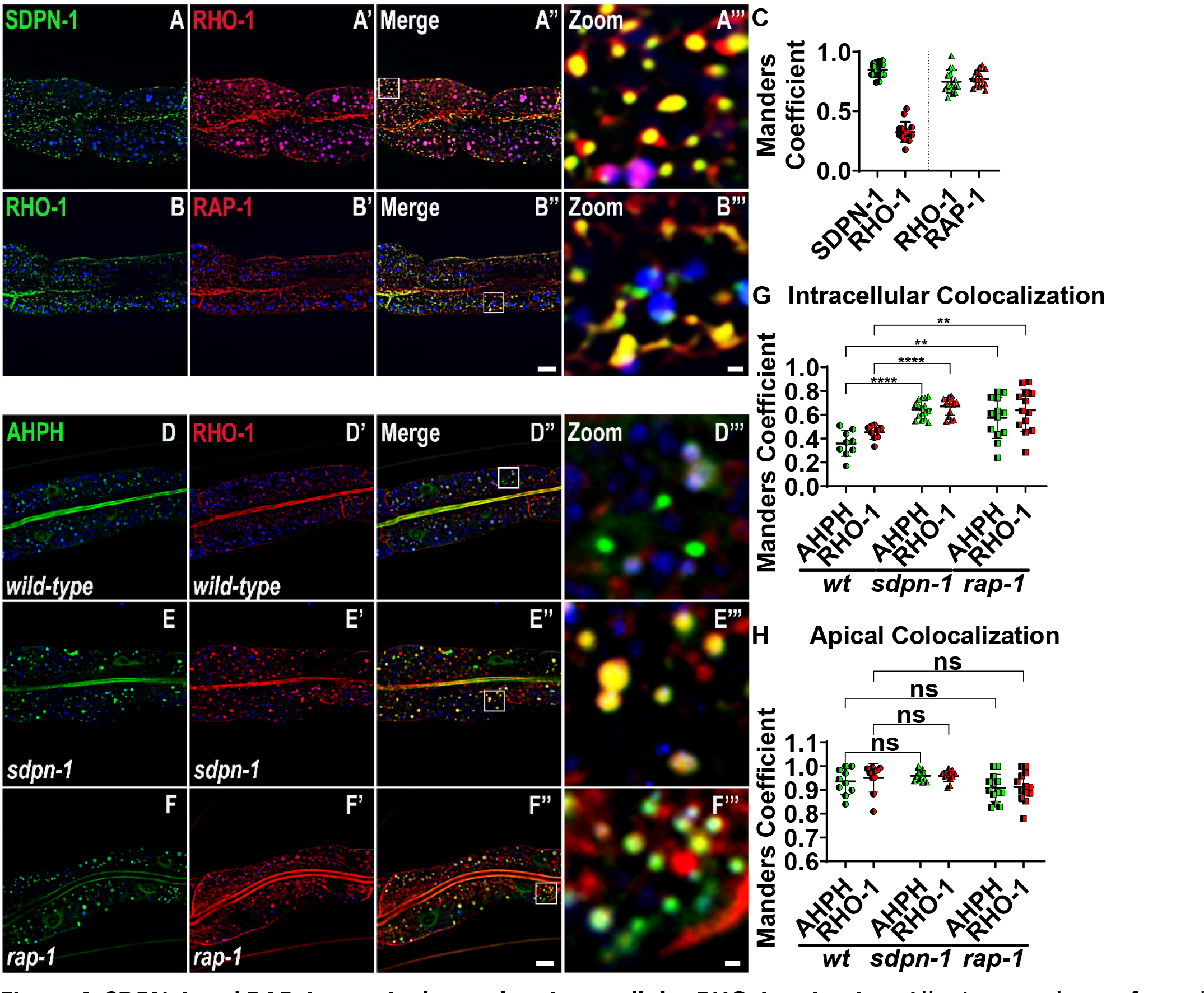
SDPN-1 and RAP-1 negatively regulate intracellular RHO-1 activation. All micrographs are from three- dimensional (3D) confocal image stacks acquired in intact living animals expressing intestine-specific GFP and RFP-tagged proteins. **(A-A’’)** tagRFP::RHO-1 colocalizes extensively with SDPN-1 positive endosomes. **(A’’’)** Magnified image from the section designated by the white rectangle in panel **(A’’)**. **(B-B’’)** GFP::RHO-1 colocalizes with RAP-1 positive endosomes. Scale bar = 10 µm. **(B’’’)** Magnified image from the section designated by the white rectangle in panel **(B’’)**. Scale bar = 1 µm. **(C)** Manders coefficient for colocalization of SDPN-1::GFP with tagRFP::RHO-1 and GFP::RHO-1 with tagRFP::RAP-1. **(D-F’’)** Imaging of active RHO-1(GTP) via colocalization of RHO-1(GTP) biosensor AHPH with tagRFP::RHO-1 in wild-type, *sdpn-1(ok1667)*, and *rap- 1(tm861)* backgrounds. Scale bar = 10 µm. **(D’’-F’’)** Magnified images from the section designated by the white rectangle in panel. Scale bar = 1 µm. **(D’’’-F’’’)**. In each image, the blue channel denotes autofluorescent lysosome-like organelles, whereas the green channel denotes GFP and the red channel denotes RFP. The green channel and red channel signal that do not overlaps with the blue channel signal are considered bona fide GFP or RFP signals respectively. **(G)** Manders coefficient for intracellular colocalization of AHPH with RHO-1 in a *wild- type*, *sdpn-1 null* and *rap-1 null* background. Loss of SDPN-1 or RAP-1 results in an increased colocalization, indicating increased activation of RHO-1 in *sdpn-1* and *rap-1* mutants. **(H)** Manders coefficient for apical colocalization of AHPH with RHO-1 in a *wild-type*, *sdpn-1 null* and *rap-1 null* background. AHPH recruitment to apical RHO-1 is not increased in *sdpn-1* and *rap-1* mutants, supporting that the key site of action of SDPN-1 on RHO-1 is the endosomes and not the apical plasma membrane. Each data point represents one animal. Error bars represent SEM: ns (non-significant), **p< 0.01, **** p< 0.0001 (Two-tailed Welch’s *t* test).

**Figure 5.**
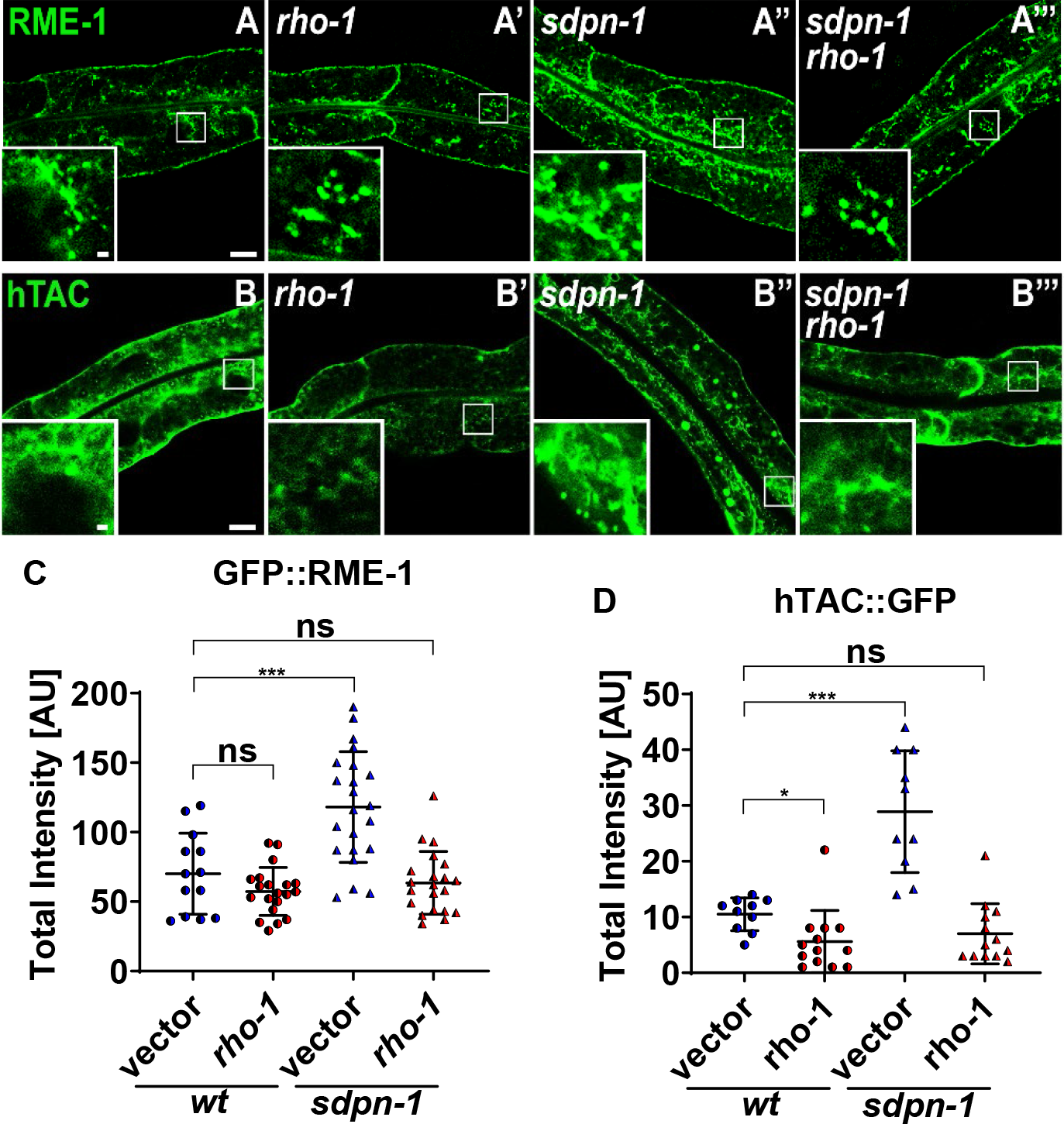
Depletion of RHO-1 suppresses *sdpn-1 null* mutant endocytic recycling phenotypes. All micrograph images were captured using confocal laser-scanning microscopy in intact living animals expressing intestine- specific GFP-tagged proteins. **(A-A’)** GFP::RME-1 positive recycling endosomes visualized in otherwise *wild-type* animals after *empty-vector RNAi* or *rho-1* RNAi. **(A’’-A’’’)** GFP::RME-1 positive recycling endosomes visualized in *sdpn-1(ok1667)* mutant animals after *empty-vector RNAi* or *rho-1* RNAi. **(B-B’)** Recycling cargo hTAC::GFP was visualized in otherwise *wild-type* animals after *empty-vector RNAi* or *rho-1* RNAi. **(B’’-B’’’)** Recycling cargo hTAC::GFP was visualized in *sdpn-1(ok1667)* mutant animals after *empty-vector RNAi* or *rho-1* RNAi. Whole image scale bar = 10 µm. The panel at the bottom left of the image is the magnified section inside the white rectangle, scale bar = 1 µm. **(C)** Quantification of intracellular total fluorescence intensity for **(A-A’’’)**. *sdpn- 1(ok1667)* mutant animals display an intracellular increase of GFP::RME-1-positive recycling endosomes. This phenotype is suppressed by *rho-1 RNAi* knockdown. **(D)** Quantification of intracellular total fluorescence intensity for **(B-B’’’)**. *sdpn-1(ok1667)* mutant animals accumulate abnormally high levels of intracellular recycling cargo hTAC::GFP. This phenotype is suppressed by *rho-1 RNAi* knockdown. Each data point represents one animal. Error bars represent SEM: ns (non-significant), *p< 0.05, ***p< 0.001 (Two-tailed Welch’s *t* test).

### SDPN-1 and RAP-1 negatively regulate RHO-1

If RHO-1 is regulated by SDPN-1 and RAP-1, we might expect to find a change in RHO-1 subcellular localization in *sdpn-1* and *rap-1* mutants. Indeed, we found an increased accumulation of GFP::RHO-1 in intracellular puncta in both mutants (Figure S4). This result was suggestive, but was not sufficient to define any changes in RHO-1 GTP-level.

To directly test for effects of *sdpn-1* and *rap-1* mutants on intestinal endosomal RHO-1 activation state, we co-expressed a RHO-1(GTP) biosensor (GFP::AHPH) (47) with tagRFP::RHO- 1, assaying for changes in GFP::AHPH recruitment to tagRFP::RHO-1 labeled endosomes in the intestine. If SDPN-1 normally acts to negatively regulate RHO-1 activation via RAP-1 signaling, we would expect *sdpn-1* and *rap-1* mutants to show increased RHO-1 activation, leading to increased GFP::AHPH recruitment. Indeed, we measured a significant increase in the colocalization of RHO-1 and its active-state biosensor on intracellular structures in *sdpn-1* or *rap-1* null mutant backgrounds, indicating abnormally high RHO-1(GTP) levels on intestinal endosomes (Fig. 4D-G). Importantly, there was no change in GFP::AHPH colocalization and tagRFP::RHO-1 on the apical plasma membrane, indicating the specificity of this regulation (Fig 4H). This data suggests a model in which a critical function of the SDPN-1/PXF-1/RAP-1 pathway is to prevent the inappropriate overactivation of RHO-1 on endosomes that could interfere with recycling function.

### RHO-1 functions downstream of SDPN-1 to regulate endocytic recycling

A corollary to this model is that reducing RHO-1 levels could be expected to compensate for elevated RHO-1 activation in *sdpn-1* mutants, leading to rescue of *sdpn-1* endocytic recycling defects. To test this hypothesis directly, we used RNAi to deplete RHO-1 in a *sdpn-1* mutant background and assayed for effects on recycling endosomes and endocytic recycling cargo. Since *rho-1* is an essential gene, we established RNAi conditions that strongly depleted RHO-1, as verified in a tagRFP::RHO-1 strain (Fig. S5), but allowed development to the adult stage typically used in our assays. Remarkably, this RNAi-mediated depletion of RHO-1 fully suppressed *sdpn-1* null mutant defects that normally result in abnormal accumulation of GFP::RME-1-labeled recycling endosomes and hTAC::GFP recycling cargo (Fig. 6, panel A-B’’’’). These results strongly support regulation of endosomal RHO-1 as a key factor in the SDPN-1- dependent regulation of endocytic recycling.

**Figure 6.**
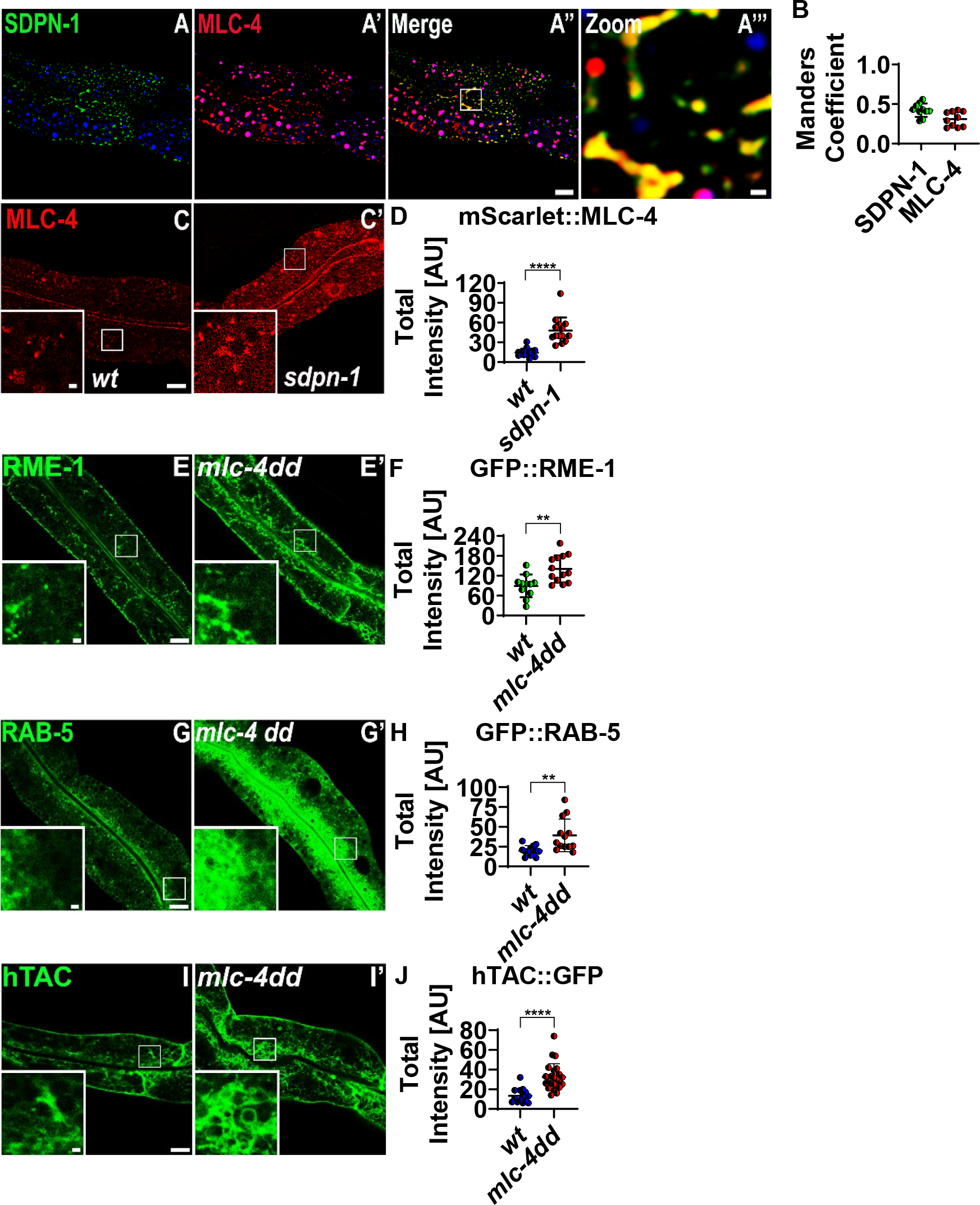
Non-muscle myosin II associates with SDPN-1-positive endosomes and modulates their function. (A-A’’) Confocal images comparing the subcellular localization of SDPN-1::GFP-positive endosomes and myosin light chain mScarlet::MLC-4 in the intestine of intact living animals. Most of SDPN-1 positive compartments are enriched in myosin light chain (MLC-4). Scale bar = 10 µm. **(A’’’)** Magnified image from the section designated by the white rectangle in panel **(A’’)**. Scale bar = 1 µm. In each image, the blue channel denotes autofluorescent lysosome-like organelles, whereas the green channel denotes GFP and the red channel denotes mScarlet. The green channel and red channel signal that does not overlaps with the blue channel signal are considered *bona fide* GFP or mScarlet signals respectively. **(B)** Manders coefficient for colocalization of SDPN-1::GFP with mScarlet::MLC-4 indicates strong enrichment for MLC-4 on endosomes. **(C-C’)** The signal of intracellular myosin light chain (mScarlet::MLC-4) positive compartments increases in a *sdpn-1(ok1667)* mutant background. **(D)** Quantification intracellular total fluorescence intensity for **(C-C’)**. **(E-H)** Animals expressing a constitutively active myosin light chain phospho-mimetic protein MLC-4DD in the intestine display increased intracellular accumulation of early and recycling endosomes. **(I-J)** Animals expressing a constitutively active myosin light chain phospho-mimetic protein MLC-4DD in the intestine display an increase in intracellular recycling cargo hTAC::GFP. Each data point represents one animal. Error bars represent SEM: **p< 0.01, **** p< 0.0001 (Two-tailed Welch’s *t* test).

### Non-muscle myosin II is a key regulator of recycling downstream of SDPN-1, RAP-1 and RHO-1

Given our results with RHO-1, we sought greater insight into which RHO-1-mediated activities influence recycling. *C. elegans* RHO-1 and mammalian RhoA are well known for regulating actomyosin contractility through non-muscle myosin II (48, 49). In support of a roll for myosin II in this pathway, we found that SDPN-1 colocalized with MLC-4, the myosin II regulatory light chain reported to function with non-muscle myosin II heavy chain isoforms NMY-1 and NMY-2 (Fig. 4, panel C-C’’’ and D). We also found that loss of SDPN-1 increased intracellular accumulation of punctate MLC-4 intracellular signal suggesting increased recruitment to endosomal membranes (50).

If regulation of myosin II contraction cycling is a key output of SDPN-1/PXF-1/RAP-1 signaling for endocytic recycling, forced activation of myosin II activity might be expected interfere with endosome function. Indeed, we found that forced activation of non-muscle myosin II in the intestine via expression of a constitutively active phosphomimetic form of MLC-4 (MLC-4(DD)) (50), increased intracellular retention of recycling cargo hTAC::GFP, and resulted in intracellular accumulation of RAB-5 and RME-1 marked endosomes (Fig. 6, panel E-H; Fig S1C’-C’’). These results support a requirement for myosin II regulation on endosomes for successful endocytic recycling.

To investigate myosin II requirements further, we assayed for effects of depletion of non-muscle myosin II isoforms NMY-1 and NMY-2. Importantly, depletion of myosin II isoform NMY- 2 caused intracellular accumulation of recycling endosome marker RME-1 and recycling cargo hTAC::GFP similar to *sdpn-1* mutants, while depletion of NMY-1 did not (Fig 7 H-J, G). By contrast, depletion of myosin II isoform NMY-1, but not depletion of NMY-2, strongly suppressed *sdpn-1* null mutant intracellular accumulation of GFP::RME-1-labeled recycling endosomes and recycling cargo hTAC::GFP, similar to the effects of RHO-1 depletion (Fig. 7).

**Figure 7.**
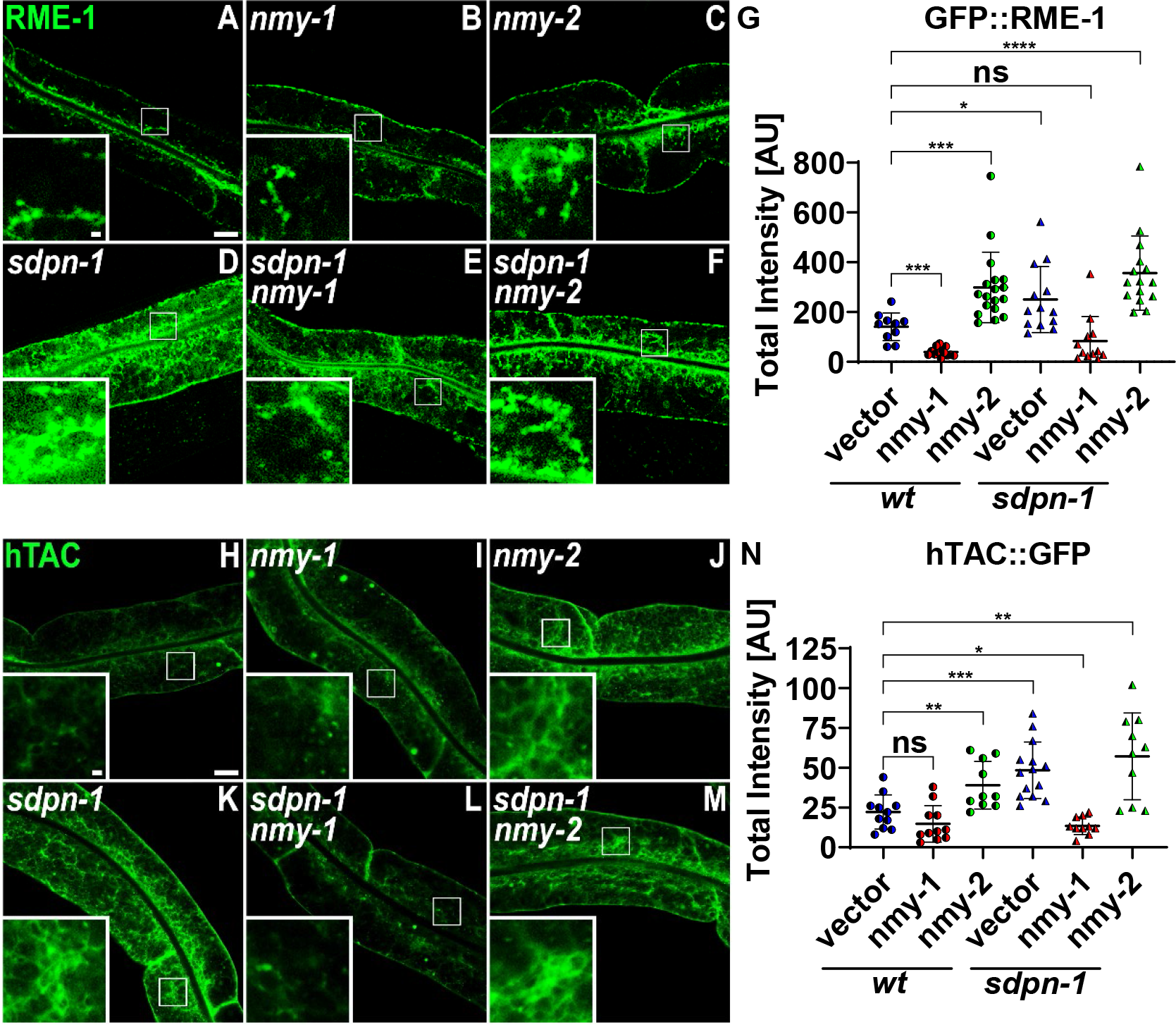
Opposing requirements for non-muscle myosin heavy chains on endocytic recycling. All micrograph images were captured using confocal laser-scanning microscopy in intact living animals expressing intestine-specific GFP-tagged proteins. **(A-C)** GFP::RME-1 positive recycling endosomes visualized in otherwise *wild-type* animals after *empty-vector* RNAi, *mny-1* RNAi or *nmy-2* RNAi. **(D-F)** GFP::RME-1 positive recycling endosomes visualized in *sdpn-1(ok1667)* mutant animals after *empty-vector* RNAi, *nmy-1* RNAi or *nmy-2* RNAi. Whole image scale bar = 10 µm. The panel at the bottom left of the image is the magnified section inside the white rectangle, scale bar = 1 µm. **(G)** Quantification of intracellular total fluorescence intensity for **(A-F)**. *sdpn-1(ok1667)* mutant animals display an increase of intracellular GFP::RME-1-positive recycling endosomes. This phenotype is suppressed by *nmy-1 RNAi* knockdown, but phenocopied by *nmy-2 RNAi* knockdown. **(H-J)** Recycling cargo hTAC::GFP visualized in otherwise *wild-type* animals after *empty-vector* RNAi, *mny-1* RNAi or *nmy-2* RNAi. **(K-M)** Recycling cargo hTAC::GFP visualized in *sdpn-1(ok1667)* mutant animals after *empty-vector* RNAi, *nmy-1* RNAi or *nmy-2* RNAi. Whole image scale bar = 10 µm. The panel at the bottom left of the image is the magnified section inside the white rectangle, scale bar = 1 µm. **(N)** Quantification of intracellular total fluorescence intensity for **(H-M)**. *sdpn-1(ok1667)* mutant animals accumulate abnormally high levels of intracellular recycling cargo hTAC::GFP. This phenotype is suppressed by *nmy-1 RNAi* knockdown, but phenocopied by *nmy-2 RNAi* knockdown. Each data point represents one animal. Error bars represent SEM: ns (non-significant), *p< 0.05, **p< 0.01, ***p< 0.001, ****p< 0.0001 (Two-tailed Welch’s *t* test).

Taken together these results strongly support a key role for myosin II regulation on endosomes in recycling function, with NMY-1 as a likely target of SDPN-1/PXF-1/RAP-1/RHO-1 regulation (Fig.8).

**Figure 8.**
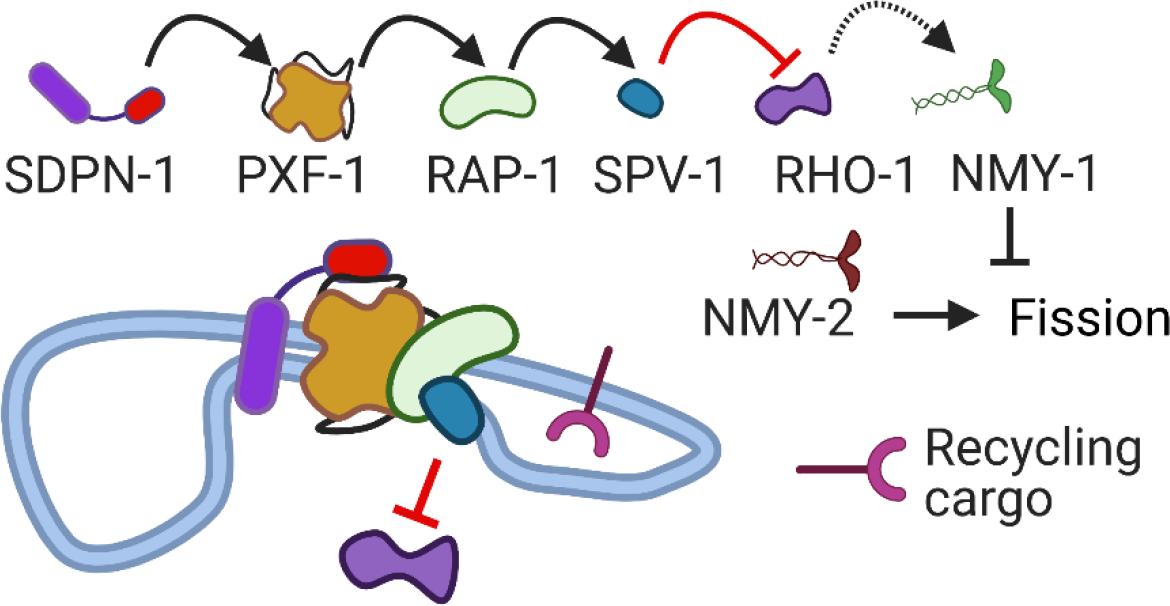
Model: SDPN-1 differentially regulates Non-Muscle Myosins via RAP-1 to RHO-1 signaling to regulate endocytic recycling. Recycling tubules pulled from sorting endosomes require non-muscle myosin NMY-2, likely for tubule fission. The F-BAR protein SDPN-1/Syndapin and its binding partner PXF-1 activate RAP-1/Rap to restrain the level of RHO-1/RhoA and non-muscle myosin NMY-1 activity on the endosome. Such negative regulation is required for successful recycling.

## Discussion

Syndapin/PACSIN proteins have long been implicated in endocytic recycling in multiple organisms, but a clear understanding of the mechanism of action of these proteins in endosomal regulation has remained unclear (Braun et al., 2005; Farmer et al., 2021; Gleason et al., 2016). Some clues are provided by the syndapin/PACSIN domain organization that is characterized by a single N-terminal F-BAR domain that links these proteins to membranes, and a single C-terminal SH3 domain that interacts with other proteins. Our new *in vivo* data reinforces the essential nature of the SH3 domain to syndapin/PACSIN function in recycling, finding that a point mutation in the SH3 domain of endogenous *sdpn-1* produced endocytic recycling defects in the *C. elegans* intestine equivalent to those we previously established for *sdpn-1* null mutants. Thus, identifying the key physiologically relevant binding partners of the SDPN-1 SH3 domain is essential to understanding how SDPN-1 controls recycling.

The *C. elegans* SH3-ome project predicted a high likelihood binding site for the SDPN-1 SH3 in RAP-GEF PXF-1, an interaction that we confirmed biochemically. We found the importance of the interaction *in vivo* to be clear, as mutation of this binding site in endogenous *pxf-1* also severely impaired endocytic recycling, causing intracellular accumulation of intestinal basolateral recycling cargo and abnormal accumulation of markers of early and basolateral recycling endosomes. The role of PXF-1 in recycling appears to match its known biochemical activity as a Rap-GEF, as our results further showed that loss of PXF-1 target RAP-1 also impaired intestinal recycling, and PXF-1 and RAP-1 are enriched on SDPN-1 labeled endosomes as would be expected if they function together. Although Rap GTPases have not been well studied in the context of recycling, mammalian Rap1 and Rap2 have been reported to be enriched on recycling endosomes, with Rap1 supporting recycling of the GABA(B) receptor in neurons, and Rap2 supporting recycling of integrin LFA-1 in T lymphocytes (28, 51). Taken together with our data in *C. elegans* we conclude that a requirement for Rap-GTPases in endocytic recycling is evolutionarily conserved. Thus, gaining insight into how Rap GTPases are regulated on recycling endosomes, and understanding the mechanisms by which Rap activation influences recycling function in *C. elegans* should be broadly applicable to other organisms, and has important implications for higher order processes known to require Rap signaling, including cell adhesion, cell migration, cell polarity, and cell fate specification.

What then is the function of RAP-1 signaling from endosomes during recycling? One known output of Rap signaling is the negative regulation of RhoA, a function first shown during neurite outgrowth, and also prominent during endothelial barrier formation (41, 52). Indeed, we found that *C. elegans* RHO-1/RhoA was highly enriched on early and basolateral recycling endosomes in the adult *C. elegans* intestine, and RHO-1 activation was increased upon loss of SDPN-1 or RAP-1 as judged by a RHO-1(GTP) biosensor, indicating that SDPN-1 and RAP-1 act as negative regulators of RHO-1 activity on endosomes. Importantly, our results further revealed RHO-1 negative regulation as a key output of SDPN-1 signaling relevant to recycling, since a reduction in RHO-1 levels by RNAi strongly suppressed the recycling defects of *sdpn-1* mutants. We note that recent work in mammalian epithelial cells found cortical recycling endosomes as a major site of RhoA activation, in this case relevant to stress fiber generation and front-rear polarity acquisition, although this work did not test for effects of RhoA activation on recycling function (53). These similarities further suggest that our findings will be relevant in a broad context across species.

We further sought to determine which output of RHO-1 signaling was most relevant to SDPN-1- mediated recycling regulation. RhoA is a well-established regulator of actomyosin contractility that functions through effectors mDia1 (a diaphanous-related formin) and ROCK (Rho-associated serine/threonine kinase), with ROCK activating non-muscle myosin II and actin dynamics (48). Our experiments found that depletion of the NMY-1 isoform of non-muscle myosin II was sufficient to significantly suppress *sdpn-1* mutant defects. These results indicate that inappropriately high levels of active myosin II, or the inability to move through complete myosin contractility cycles, explains at least part of the effects of *sdpn-1* mutants on recycling (Gleason et al., 2016; Yan et al., 2021). This interpretation is supported by our results showing that constitutive activation of myosin II by expression of a myosin light chain phosphomimetic also impairs recycling function, and our findings that myosin light chain is enriched on SDPN-1 positive endosomes.

Surprisingly, we found the opposite phenotype for non-muscle myosin isoform NMY-2. Depletion of NMY-2 caused intracellular accumulation of recycling cargo, indicating a positive role for NMY-2 in recycling. Interestingly, previous work on NMY-1 and NMY-2 documented opposing effects in the regulation of the contractile ring channels of the *C. elegans* gonad (22). Mathematical modeling in that work suggested that these opposing effects could be explained by NMY-1 and NMY-2 contractile fibers exerting forces in opposing directions, one normal to the ring, and the other tangential (22). If these principles apply to the recycling endosome case, NMY-2 derived forces might increase tension on endosomal tubules to promote their fission, while NMY-1 contraction in an opposing direction might reduce tubule tension and fission.

Normal recycling tubule release might require cycles of increased and decreased tension, or forces regulating recycling rates might be modulated in response to changes in cellular requirements, such as changes in cargo load or metabolism.

Interestingly, three putative Rho effectors, formin CYK-1/mDia, LET-502/ROCK, and RTKN- 1/Rhotekin, are known to function in basolateral recycling in the *C. elegans* intestine, although current models posit that these proteins function independently of RHO-1 in recycling (11, 54, 55). Branched actin formed by Arp2/3 is traditionally considered the relevant F-actin type associated with endosomes, but recycling may require a balance of branched and linear actin for proper function. Actin tracks produced by CYK-1 could be substrates upon which cycles of myosin II contractility would act, potentially producing cycles of actomyosin-driven endosomal membrane tension that could promote cycles of tubular cargo carrier fission.

We note that non-muscle myosin II was previously shown to promote mitochondrial fission and could contribute to endosomal fission during recycling in a similar way (56–59). Myosin II, Spire1C, and endoplasmic reticulum-bound formin INF2 have been proposed to create actomyosin tension that deforms the outer mitochondrial membrane during “prefission”, promoting curvature-sensing-based recruitment of dynamin-related protein Drp1 and other factors that complete mitochondrial constriction (59). Further work will be required to determine if a similar process is operative at the early endosome – recycling endosome interface, and if there is a relationship between SDPN-1/RAP-1/RHO-1/NMY-1 and previously identified functions for LET-502/RTKN-1/CYK-1 during recycling.

Syndapin/PACSIN proteins are often thought of as acting directly in endosomal tubule generation or tubule severing, since isolated BAR domains from several of these proteins are able to generate tubules from liposomes *in vitro*, and in some cases enhance endosomal tubulation when overexpressed *in vivo*. However, it remains unclear if an F-BAR-driven constricting/tubulating activity is a key output of syndapin/PACSINs that promotes recycling function. An alternate hypothesis is that the key function of the F-BAR domain in syndapin/PACSINs is to provide membrane binding and curvature-sensing that brings syndapin/PACSIN to endosomal tubules elaborated by other means, such as cytoskeletal pushing and pulling forces. In the *C. elegans* intestine and human HeLa cells RAB-10/Rab10 is essential for the extensive tubulation found in the recycling endosome network (10, 60–64). In the *C. elegans* case microfilament and microtubule integrity is required for network structure, and RAB-10 is known to regulate association of endosomal membranes with actin filaments via the EHBP-1 protein (60, 63, 64). In the HeLa case Rab10-mediated regulation of Kif13/kinesin pulling forces have also been implicated (61).

Our new data clearly points to an essential role for Syndapin in signaling via Rap and Rho to control myosin II on endosomes (Fig.8), but we cannot rule out an additional mechanical contribution of SDPN-1 to endosomal function via direct membrane bending. However, the high efficiency of *sdpn-1* mutant recycling phenotype suppression by *rho-1* depletion suggests that RHO-1 regulation is the main output relevant to recycling. We also note that in some systems syndapin/PACSIN proteins associate with other trafficking regulators such as MICAL-L1, OCRL1, Dynamin, and EHD1/RME-1 that could promote tubulation and/or severing (62). It will be of great interest to revisit the question of relative contributions to the tubule-based recycling mechanism as new genetic and cell biological tools evolve, allowing more complete, acute, and fine-tuned control of experimental conditions, as we further elaborate our understanding of this complex system.

## Acknowledgments

We would like to thank the Grant lab for critical comments and suggestions. We thank Peter Schweinsberg and Ge Bai for technical assistance in making plasmid clones and biolistic transgenic lines, and Helen Ushakov for expert microinjection. We thank Adriana Dawes for *nmy-1* and *nmy-2* RNAi clones, Michel Labouesse for *mlc-4* wild-type and phosphomimetic DNA clones, and David Reiner for *rap-1* mutant strains. We also thank Michael Pierce, Noriko Goldsmith and Jessica Shivas for help with confocal microscopy, and Gary Bader for SH3 weighted match analysis. This work was supported by NIH Grant 5R01GM135326 to B.D.G.

---

**Figure S1.**
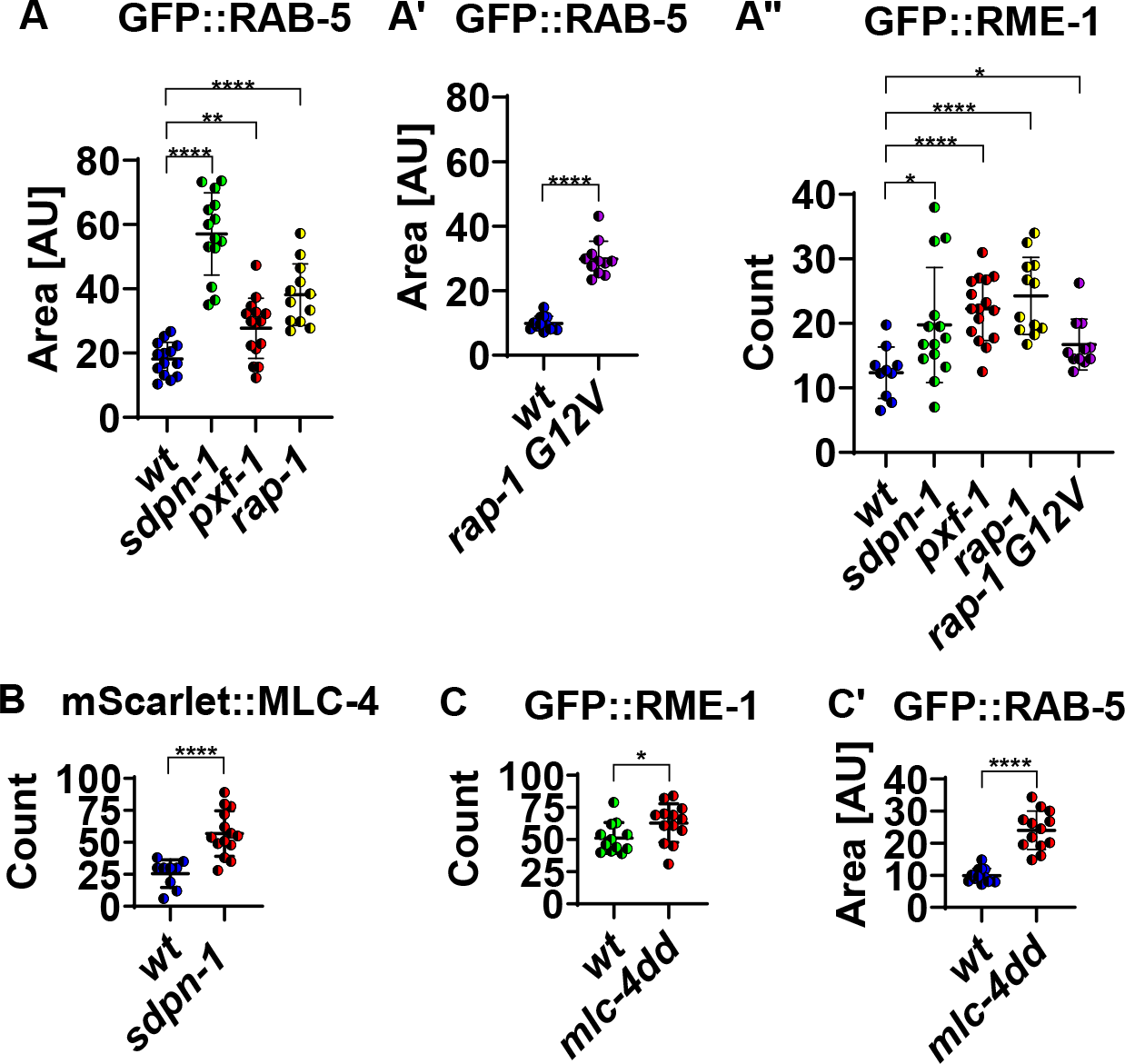
Additional quantification related to main. figures 2**, 5, and 6. (A-A’’)** Increased intracellular area labeled by GFP::RAB-5 and increased number of intracellular recycling compartments in interaction-defective mutants *sdpn-1(pw33)* and *pxf-1(pw34)*, as well as null *rap-1(tm861)* and activated *rap-1(re180)* (*rap-1 G12V*) mutants (related to Main Fig. 2A-C’’’’). **(B)** Quantification of the number of intracellular mScarlet::MLC-4 regulatory myosin light chain puncta (related to Main Fig. 6C-C’). **(C-C’)** Quantification of GFP::RME-1 labeled recycling compartments and intracellular area positive for early endosome marker GFP::RAB-5 (related to main Fig. 6E-H). Error bars represent SEM: *p< 0.05, **p< 0.01, **** p< 0.0001 (Two-tailed Welch’s *t* test).

**Figure S2.**
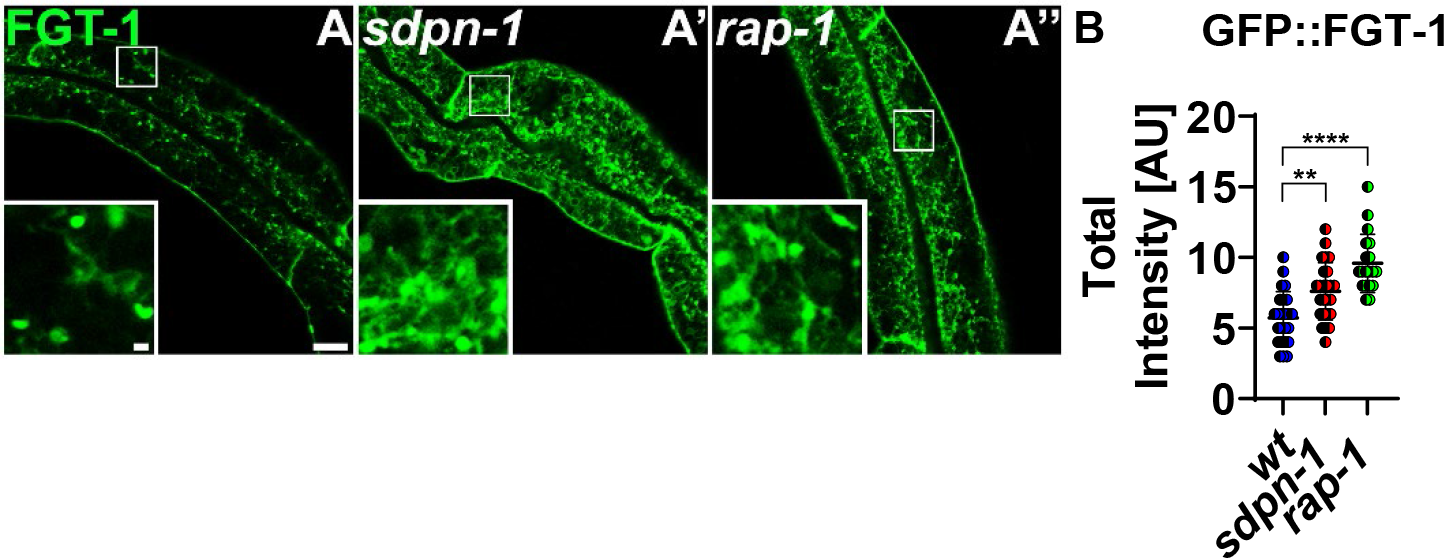
Intracellular accumulation of glucose transporter FGT-1 in *sdpn-1* and *rap-1 null* mutants. All micrograph images were obtained by confocal laser-scanning microscopy in intact living animals expressing intestine-specific GFP-tagged protein. **(A-A’’)**. Representative images are shown for FGT-1::GFP in wild-type, *sdpn-1(ok1667),* and *rap-1(tm861)* mutants. Whole image scale bar = 10 µm. The panel at the bottom left of the image is the magnified section inside the white rectangle, scale bar = 1 µm. **(B)** Quantification of total intracellular fluorescence intensity. There is a significant increase in intracellular FGT-1::GFP accumulation in a *sdpn-1(0)* and *rap-1(0)* mutants compared to *wild-type*. Each data point represents one animal. Error bars represent SEM: ** p<0.01, **** p< 0.0001 (Two-tailed Welch’s *t* test).

**Figure S3.**
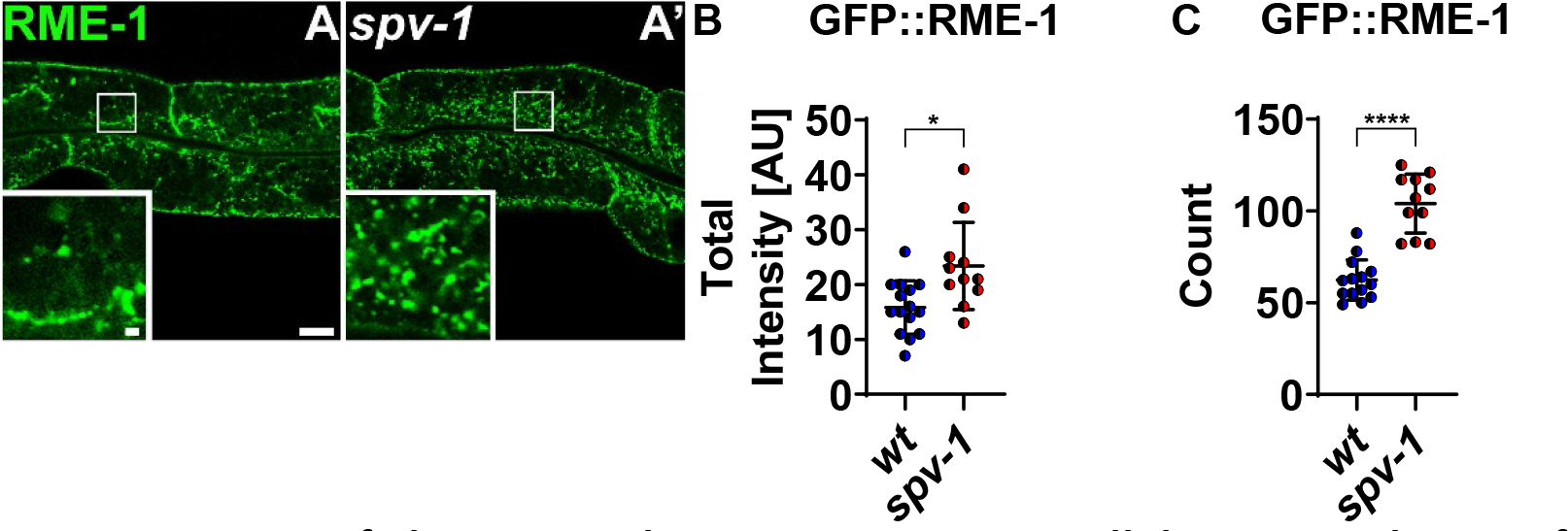
Loss of the SPV-1 Rho-GAP causes intracellular accumulation of RME-1-positive recycling endosomes. All micrograph images were obtained by confocal laser-scanning microscopy in intact living animals expressing intestine-specific GFP-tagged protein. **(A-A’)** Representative images are shown for GFP::RME-1 labeled recycling endosome in wild-type and *spv-1(ok1498)* mutant animals. Whole image scale bar = 10 µm. The panel at the bottom left of the image is the magnified section inside the white rectangle, scale bar = 1 µm. **(B)** Quantification of total intracellular fluorescence intensity of recycling endosomes. **(C)** Quantification of the number of intracellular recycling endosomes. There is a significant increase in RME-1-positive recycling endosomes in *spv-1* mutants. Each data point represents one animal. Error bars represent SEM: * p< 0.05, **** p< 0.0001 (Two-tailed Welch’s *t* test).

**Figure S4.**
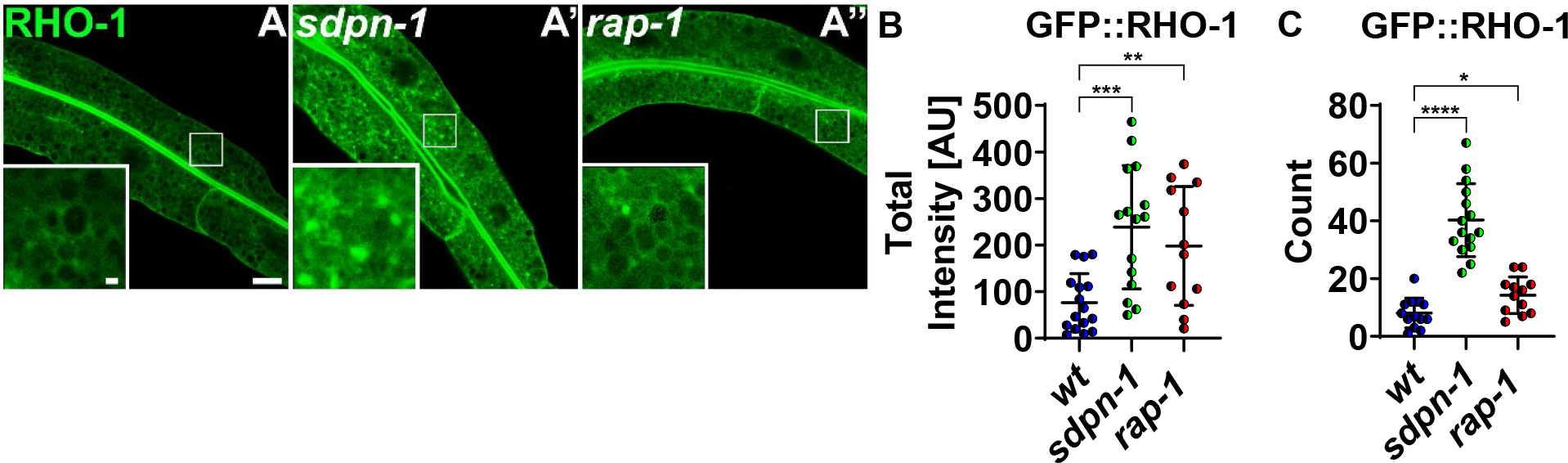
Intracellular RHO-1 signal increases in *sdpn-1* and *rap-1 null* mutants. All micrograph images were obtained via confocal laser-scanning microscopy in intact living animals expressing intestine-specific GFP-tagged protein. **(A-A’’)** Whole image scale bar = 10 µm. The panel at the bottom left of the image is the magnified section inside the white rectangle, scale bar = 1 µm. **(B)** Quantification of total intracellular fluorescence intensity of RHO-1. **(C)** Quantification of the number of intracellular RHO-1 puncta. Intracellular puncta signal for GFP::RHO-1 increases significantly in *sdpn-1(ok1667)* and *rap-1(tm861)* mutants. Each data point represents one animal. Error bars represent SEM: * p< 0.05, ** p< 0.01, *** p< 0.001, **** p< 0.0001 (Two-tailed Welch’s *t* test).

**Figure S5.**
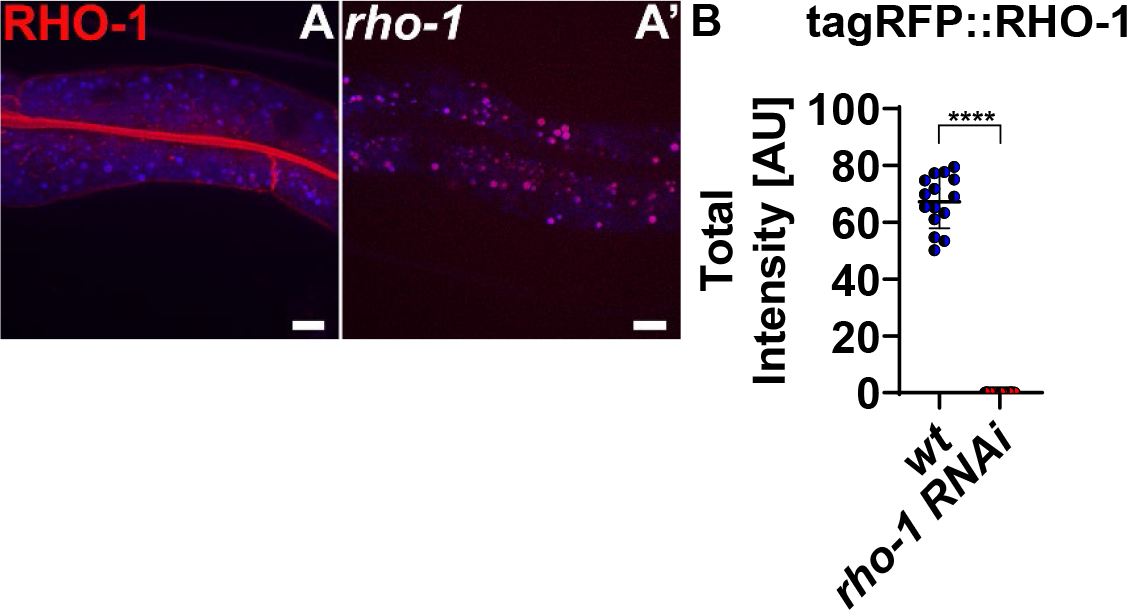
Intestinal RHO-1 was depleted by RHO-1 RNAi. All micrographs are from confocal image stacks acquired in intact living animals expressing intestine-specific RFP-tagged protein. **(A)** tagRFP::RHO-1 expression on a wild-type background. **(A’)** tagRFP::RHO-1 is depleted by rho- 1 RNAi. Scale bar = 10 µm. In each image, the blue channel denotes autofluorescent lysosome- like organelles, whereas the red channel denotes RFP. The red channel signal that does not overlap with the blue channel signal is considered bona fide RFP signal respectively. **(B)** Quantification of total fluorescence intensity. Each data point represents one animal. Error bars represent SEM: **** p< 0.0001 (Two-tailed Welch’s *t* test).

## References

1. G. J. Doherty, H. T. McMahon, Mechanisms of endocytosis. Annu Rev Biochem 78, 857–902 (2009).

2. L. Maldonado-Báez, C. Williamson, J. G. Donaldson, Clathrin-independent endocytosis: a cargo- centric view. Exp Cell Res 319, 2759–2769 (2013).

3. J. A. Solinger, A. Spang, Sorting of cargo in the tubular endosomal network. Bioessays 44, e2200158 (2022).

4. B. D. Grant, J. G. Donaldson, Pathways and mechanisms of endocytic recycling. Nat Rev Mol Cell Biol 10, 597–608 (2009).

5. J. D. McGhee, The C. elegans intestine. WormBook 10.1895/wormbook.1.133.1, 1-36 (2007).

6. B. Leung, G. J. Hermann, J. R. Priess, Organogenesis of the Caenorhabditis elegans intestine. Dev Biol 216, 114–134 (1999).

7. B. Grant et al., Evidence that RME-1, a conserved C. elegans EH-domain protein, functions in endocytic recycling. Nat Cell Biol 3, 573–579 (2001).

8. K. Sato, A. Norris, M. Sato, B. D. Grant, C. elegans as a model for membrane traffic. WormBook 10.1895/wormbook.1.77.2, 1–47 (2014).

9. A. M. Gleason, K. C. Nguyen, D. H. Hall, B. D. Grant, Syndapin/SDPN-1 is required for endocytic recycling and endosomal actin association in the C. elegans intestine. Mol Biol Cell 27, 3746–3756 (2016).

10. C. C. Chen et al., RAB-10 is required for endocytic recycling in the Caenorhabditis elegans intestine. Mol Biol Cell 17, 1286–1297 (2006).

11. Y. Yan et al., RTKN-1/Rhotekin shields endosome-associated F-actin from disassembly to ensure endocytic recycling. J Cell Biol 220 (2021).

12. E. Casal et al., The crystal structure of the BAR domain from human Bin1/amphiphysin II and its implications for molecular recognition. Biochemistry 45, 12917–12928 (2006).

13. A. Braun et al., EHD proteins associate with syndapin I and II and such interactions play a crucial role in endosomal recycling. Mol Biol Cell 16, 3642–3658 (2005).

14. A. Quan, P. J. Robinson, Syndapin--a membrane remodelling and endocytic F-BAR protein. Febs j 280, 5198–5212 (2013).

15. J. Kovár, Hybridoma cultivation in defined serum-free media: growth-supporting substances. II. Insulin, other hormones, and growth factors. Folia Biol (Praha) 32, 304–310 (1986).

16. X. Xin et al., SH3 interactome conserves general function over specific form. Mol Syst Biol 9, 652 (2013).

17. W. Pellis-van Berkel et al., Requirement of the Caenorhabditis elegans RapGEF pxf-1 and rap-1 for epithelial integrity. Mol Biol Cell 16, 106–116 (2005).

18. M. R. Kooistra, N. Dubé, J. L. Bos, Rap1: a key regulator in cell-cell junction formation. J Cell Sci 120, 17–22 (2007).

19. C. Frøkjær-Jensen et al., Random and targeted transgene insertion in Caenorhabditis elegans using a modified Mos1 transposon. Nat Methods 11, 529–534 (2014).

20. L. Timmons, A. Fire, Specific interference by ingested dsRNA. Nature 395, 854 (1998).

21. R. S. Kamath, J. Ahringer, Genome-wide RNAi screening in Caenorhabditis elegans. Methods 30, 313–321 (2003).

22. V. C. Coffman, T. M. Kachur, D. B. Pilgrim, A. T. Dawes, Antagonistic Behaviors of NMY-1 and NMY-2 Maintain Ring Channels in the C. elegans Gonad. Biophys J 111, 2202–2213 (2016).

23. A. Paix, A. Folkmann, G. Seydoux, Precision genome editing using CRISPR-Cas9 and linear repair templates in C. elegans. Methods 121-122, 86-93 (2017).

24. G. Huang et al., Improved CRISPR/Cas9 knock-in efficiency via the self-excising cassette (SEC) selection method in C. elegans. MicroPubl Biol 2021 (2021).

25. M. L. Schwartz, M. W. Davis, M. S. Rich, E. M. Jorgensen, High-efficiency CRISPR gene editing in C. elegans using Cas9 integrated into the genome. PLoS Genet 17, e1009755 (2021).

26. Q. Wang et al., Molecular mechanism of membrane constriction and tubulation mediated by the F-BAR protein Pacsin/Syndapin. Proc Natl Acad Sci U S A 106, 12700–12705 (2009).

27. H. Meng et al., PACSIN 2 represses cellular migration through direct association with cyclin D1 but not its alternate splice form cyclin D1b. Cell Cycle 10, 73–81 (2011).

28. P. Stanley, S. Tooze, N. Hogg, A role for Rap2 in recycling the extended conformation of LFA-1 during T cell migration. Biol Open 1, 1161–1168 (2012).

29. F. Balzac et al., E-cadherin endocytosis regulates the activity of Rap1: a traffic light GTPase at the crossroads between cadherin and integrin function. J Cell Sci 118, 4765–4783 (2005).

30. L. Li et al., A unique interplay between Rap1 and E-cadherin in the endocytic pathway regulates self-renewal of human embryonic stem cells. Stem Cells 28, 247–257 (2010).

31. Y. Liao et al., RA-GEF, a novel Rap1A guanine nucleotide exchange factor containing a Ras/Rap1A-associating domain, is conserved between nematode and humans. J Biol Chem 274, 37815–37820 (1999).

32. H. B. Kuiperij et al., Characterisation of PDZ-GEFs, a family of guanine nucleotide exchange factors specific for Rap1 and Rap2. Biochim Biophys Acta 1593, 141–149 (2003).

33. B. Qualmann, J. Roos, P. J. DiGregorio, R. B. Kelly, Syndapin I, a synaptic dynamin-binding protein that associates with the neural Wiskott-Aldrich syndrome protein. Mol Biol Cell 10, 501–513 (1999).

34. A. Shi et al., RAB-10-GTPase-mediated regulation of endosomal phosphatidylinositol-4,5- bisphosphate. Proc Natl Acad Sci U S A 109, E2306–2315 (2012).

35. R. J. Gleason, A. M. Akintobi, B. D. Grant, R. W. Padgett, BMP signaling requires retromer- dependent recycling of the type I receptor. Proc Natl Acad Sci U S A 111, 2578–2583 (2014).

36. E. W. Frische et al., RAP-1 and the RAL-1/exocyst pathway coordinate hypodermal cell organization in Caenorhabditis elegans. Embo j 26, 5083–5092 (2007).

37. J. F. Rebhun, A. F. Castro, L. A. Quilliam, Identification of guanine nucleotide exchange factors (GEFs) for the Rap1 GTPase. Regulation of MR-GEF by M-Ras-GTP interaction. J Biol Chem 275, 34901–34908 (2000).

38. N. R. Rasmussen, D. J. Dickinson, D. J. Reiner, Ras-Dependent Cell Fate Decisions Are Reinforced by the RAP-1 Small GTPase in Caenorhabditiselegans. Genetics 210, 1339–1354 (2018).

39. Y. Feng, B. G. Williams, F. Koumanov, A. J. Wolstenholme, G. D. Holman, FGT-1 is the major glucose transporter in C. elegans and is central to aging pathways. Biochem J 456, 219–229 (2013).

40. V. Pizon, M. Desjardins, C. Bucci, R. G. Parton, M. Zerial, Association of Rap1a and Rap1b proteins with late endocytic/phagocytic compartments and Rap2a with the Golgi complex. J Cell Sci 107 **( Pt** **6**), 1661–1670 (1994).

41. A. Post, W. J. Pannekoek, B. Ponsioen, M. J. Vliem, J. L. Bos, Rap1 Spatially Controls ArhGAP29 To Inhibit Rho Signaling during Endothelial Barrier Regulation. Mol Cell Biol 35, 2495–2502 (2015).

42. A. Post et al., Rasip1 mediates Rap1 regulation of Rho in endothelial barrier function through ArhGAP29. Proc Natl Acad Sci U S A 110, 11427–11432 (2013).

43. G. A. Smolen, et al., A Rap GTPase interactor, RADIL, mediates migration of neural crest precursors. Genes Dev 21, 2131–2136 (2007).

44. C. W. Wilson et al., Rasip1 regulates vertebrate vascular endothelial junction stability through Epac1-Rap1 signaling. Blood 122, 3678–3690 (2013).

45. Z. Meng et al., RAP2 mediates mechanoresponses of the Hippo pathway. Nature 560, 655–660 (2018).

46. B. Saenz-Narciso, E. Gomez-Orte, A. Zheleva, I. Gastaca, J. Cabello, Control of developmental networks by Rac/Rho small GTPases: How cytoskeletal changes during embryogenesis are orchestrated. Bioessays 38, 1246–1254 (2016).

47. Y. C. Tse et al., RhoA activation during polarization and cytokinesis of the early Caenorhabditis elegans embryo is differentially dependent on NOP-1 and CYK-4. Mol Biol Cell 23, 4020–4031 (2012).

48. M. Chircop, Rho GTPases as regulators of mitosis and cytokinesis in mammalian cells. Small GTPases 5 (2014).

49. B. Sáenz-Narciso, E. Gómez-Orte, A. Zheleva, I. Gastaca, J. Cabello, Control of developmental networks by Rac/Rho small GTPases: How cytoskeletal changes during embryogenesis are orchestrated. Bioessays 38, 1246–1254 (2016).

50. C. Gally et al., Myosin II regulation during C. elegans embryonic elongation: LET-502/ROCK, MRCK-1 and PAK-1, three kinases with different roles. Development 136, 3109–3119 (2009).

51. Z. Zhang et al., GABAB receptor promotes its own surface expression by recruiting a Rap1- dependent signaling cascade. J Cell Sci 128, 2302–2313 (2015).

52. T. Yamada, T. Sakisaka, S. Hisata, T. Baba, Y. Takai, RA-RhoGAP, Rap-activated Rho GTPase- activating protein implicated in neurite outgrowth through Rho. J Biol Chem 280, 33026–33034 (2005).

53. C. Gaston et al., EpCAM promotes endosomal modulation of the cortical RhoA zone for epithelial organization. Nat Commun 12, 2226 (2021).

54. T. Gong et al., PTRN-1/CAMSAP promotes CYK-1/formin-dependent actin polymerization during endocytic recycling. Embo j 37 (2018).

55. W. Zhang et al., LET-502/ROCK Regulates Endocytic Recycling by Promoting Activation of RAB-5 in a Distinct Subpopulation of Sorting Endosomes. Cell Rep 32, 108173 (2020).

56. F. Korobova, T. J. Gauvin, H. N. Higgs, A role for myosin II in mammalian mitochondrial fission. Curr Biol 24, 409–414 (2014).

57. U. Manor et al., A mitochondria-anchored isoform of the actin-nucleating spire protein regulates mitochondrial division. Elife 4 (2015).

58. A. L. Hatch, P. S. Gurel, H. N. Higgs, Novel roles for actin in mitochondrial fission. J Cell Sci 127, 4549–4560 (2014).

59. C. Yang, T. M. Svitkina, Ultrastructure and dynamics of the actin-myosin II cytoskeleton during mitochondrial fission. Nat Cell Biol 21, 603–613 (2019).

60. A. Shi et al., EHBP-1 functions with RAB-10 during endocytic recycling in Caenorhabditis elegans. Mol Biol Cell 21, 2930–2943 (2010).

61. K. Etoh, M. Fukuda, Rab10 regulates tubular endosome formation through KIF13A and KIF13B motors. J Cell Sci 132 (2019).

62. T. Farmer et al., Defining the protein and lipid constituents of tubular recycling endosomes. J Biol Chem 296, 100190 (2021).

63. P. Wang et al., RAB-10 Promotes EHBP-1 Bridging of Filamentous Actin and Tubular Recycling Endosomes. PLoS Genet 12, e1006093 (2016).

64. S. Chen et al., SEC-10 and RAB-10 coordinate basolateral recycling of clathrin-independent cargo through endosomal tubules in Caenorhabditis elegans. Proc Natl Acad Sci U S A 111, 15432–15437 (2014).

